# DIP2 is a unique regulator of diacylglycerol lipid homeostasis in eukaryotes

**DOI:** 10.1101/2022.02.07.479356

**Authors:** Sudipta Mondal, Priyadarshan Kinatukara, Shubham Singh, Sakshi Shambhavi, Gajanan S. Patil, Noopur Dubey, Salam Herojeet Singh, Biswajit Pal, P Chandra Shekar, Siddhesh S. Kamat, Rajan Sankaranarayanan

**Affiliations:** CSIR-Centre for Cellular and Molecular Biology, Habsiguda, Hyderabad-50007, India.; Department of Biology, Indian Institute of Science Education and Research (IISER) Pune, Dr. Homi Bhabha Road, Pashan, Pune, Maharashtra 411008, India.; Academy of Scientific and Innovative Research (AcSIR), Ghaziabad-201002, India.

**Keywords:** DIP2, Diacylglycerol, Fatty acyl-AMP ligase, Lipid homeostasis, Opisthokonta, Triacylglycerol

## Abstract

Chain-length specific subsets of diacylglycerol (DAG) lipids are proposed to regulate differential physiological responses ranging from signal transduction to modulation of the membrane properties. However, the mechanism or molecular players regulating the subsets of DAG species remains unknown. Here, we uncover the role of a conserved eukaryotic protein family, DISCO-interacting protein 2 (DIP2) as a homeostatic regulator of a chemically distinct subset of DAGs using yeast, fly and mouse models. Genetic and chemical screens along with lipidomics analysis in yeast reveal that DIP2 prevents the toxic accumulation of specific DAGs in the logarithmic growth phase, which otherwise leads to endoplasmic reticulum stress. We also show that the fatty acyl-AMP ligase-like domains of DIP2 are essential for the redirection of the flux of DAG subspecies to storage lipid, triacylglycerols. Such modulation of selective DAG abundance by DIP2 is found to be crucial for optimal vacuole-membrane fusion and consequently osmoadaptation in yeast. Thus, the study illuminates an unprecedented DAG metabolism route and provides new insights on how cell fine-tunes DAG subspecies for cellular homeostasis and environmental adaptation.

## Introduction

Eukaryotic cells are functionally compartmentalized by membranes and the integrity of these compartments requires non-uniform distribution of different lipids (van Meer et al., 2008). The physicochemical properties of different lipids are attributed to the bilayer formation, regulation of membrane protein activity, membrane curvature and fusion-fission processes (Bigay & Antonny, 2012; Harayama & Riezman, 2018). DAG is one such crucial bioactive lipid functioning at the crossroads of cell signalling and core metabolic pathways including membrane biogenesis and fat storage. Intriguingly, a growing body of evidence suggests that chemically defined subsets of DAGs, with specific fatty acid (FA) tails, determine the fate of fundamental cellular processes (Marignani et al., 1996; Schuhmacher et al., 2020; Ware et al., 2020). Therefore, precise regulation of DAG subspecies through selective formation and degradation is crucial while the identity of molecular players and their mechanisms remain unexplored.

A key step of lipid metabolism is the incorporation of FA of different chain lengths and unsaturation in lipids during biosynthesis or remodelling (Grevengoed et al., 2014; Watkins, 1997). In either case, FA is activated via adenylation reaction using ATP molecules by Fatty acyl-CoA ligase (FACLs), a member of the ANL superfamily of enzymes (Gulick, 2009). However, a recently identified homolog of FACLs, named fatty acyl-AMP ligases (FAALs) also activate FA but redirect them to 4’-phosphopantetheine coupled to acyl-carrier protein (ACP) in bacteria. The complex lipids of Mycobacteria, lipopeptides in several bacteria are the classical examples of such metabolic diversions created by FAALs, where they crosstalk with polyketide synthases (PKS) and nonribosomal peptide synthetases (NRPS) (Arora et al., 2009; Goyal et al., 2012; Trivedi et al., 2004). We have recently identified a distant ortholog of FAALs as a part of a conserved three-domain protein called DIP2 across the eukaryotic supergroup Opisthokonta (Fungi and Animals) (Patil et al., 2021). However, paradoxically, most of the opisthokonts have lost PKS/NRPS gene cluster, suggesting the emergence of possible alternate functions of these FAAL orthologs. Recently, a loss of virulence has been reported when FAAL-like domain-containing protein, CPS1, a DIP2 ortholog, is mutated in either plant pathogenic fungi, which causes rice blast and wheat head scab, or human pathogenic fungus, that cause valley fever (Lu et al., 2003; Narra et al., 2016; Wang et al., 2016). Furthermore, DIP2 has been shown to be crucial for axonal branching, neurite sprouting and dendritic spine morphogenesis in *Drosophila*, *Caenorhabditis elegans* and mice, respectively (Ma et al., 2019; Nitta et al., 2017; Noblett et al., 2019). Three paralogs of DIP2 in humans have been implicated as a potential risk factor for neurodevelopmental disorders like autism spectrum disorders (ASD) and other diseases (**Supplementary Table 1**). Despite these striking physiological consequences, our understanding of how the loss of DIP2 contributes to pathogenesis is limited by the lack of information on its cellular function.

Here, we report the previously unrecognized function for the unique FAAL-like domain (FLD) containing protein, DIP2, which ensures a metabolic redirection of a defined subset of DAGs towards triacylglycerol (TAG) lipids in yeast. We provide evidence that the DIP2-mediated DAG metabolism route is conserved in *Drosophila* and mice. The mutational study suggests that the canonical enzymatic function of FLD, i.e., fatty acid activation via adenylation, is crucial for the function of DIP2 in DAG to TAG conversion. Aberrations in this metabolic flux in the absence of DIP2 trigger a homeostatic signal called unfolded protein response (UPR) pathway leading to ER stress. Using loss and gain of function experiments, we reveal the role of DIP2-mediated DAG regulation in osmoadaptation by facilitating vacuole fusion-fission homeostasis in yeast. Overall, the work presents the discovery of a key regulator of DAG subspecies in yeast and assigns the physiological role of overlooked DAG subspecies in fundamental cellular processes, that will strengthen the emerging paradigm of lipid-mediated functional diversity.

## Results

### DIP2 is a conserved and noncanonical player in DAG metabolism across Opisthokonta

Using a phylogenetic approach, we identified that FLDs underwent two independent events of duplication (Fig. 1A). The prokaryotic FAALs were recruited early in eukaryotes along with PKS/NRPS systems in several Protozoa, Bikonta (mainly plants and algae), and Fungi. Subsequently, a highly diverged FLD went through the first duplication to produce a tandemly fused FLD-didomain in all opisthokonts (except Basidiomycota fungi). The second duplication event occurred in the vertebrates resulting in multiple paralogs of DIP2 (Fig. 1A). Though FLDs cluster with prokaryotic and plant FAALs, they segregate as sub-clusters that are specific to FLD1 and FLD2 (Fig. 1B and C). Further analyses revealed that both the FLDs retain the ancestral motifs of the ANL superfamily (Gulick, 2009) while gaining exclusive variations in several motifs, unique to each FLDs (**Supplementary Fig. 1**). The divergence of Opisthokonta FLDs from prokaryotic FAALs can also be explained on the account of the low sequence identity (∼17-21%) they share. For instance, Cmr2 (YOR093c), the yeast ortholog of DIP2 (hereafter referred to as ScDIP2), shares an average sequence identity of ∼19% (∼ 16-27 %) with other DIP2 proteins. The tandem FLDs, FLD1 and FLD2, share an average sequence identity of ∼19% and ∼22% with respective FLDs from other DIP2 and ∼16% for both the FLDs with the representative bacterial FAALs. Taken together, the data suggest that the duplication and divergence of FLDs have resulted in the emergence of DIP2 as a distinct gene family with possible new functions.

**Fig. 1.**
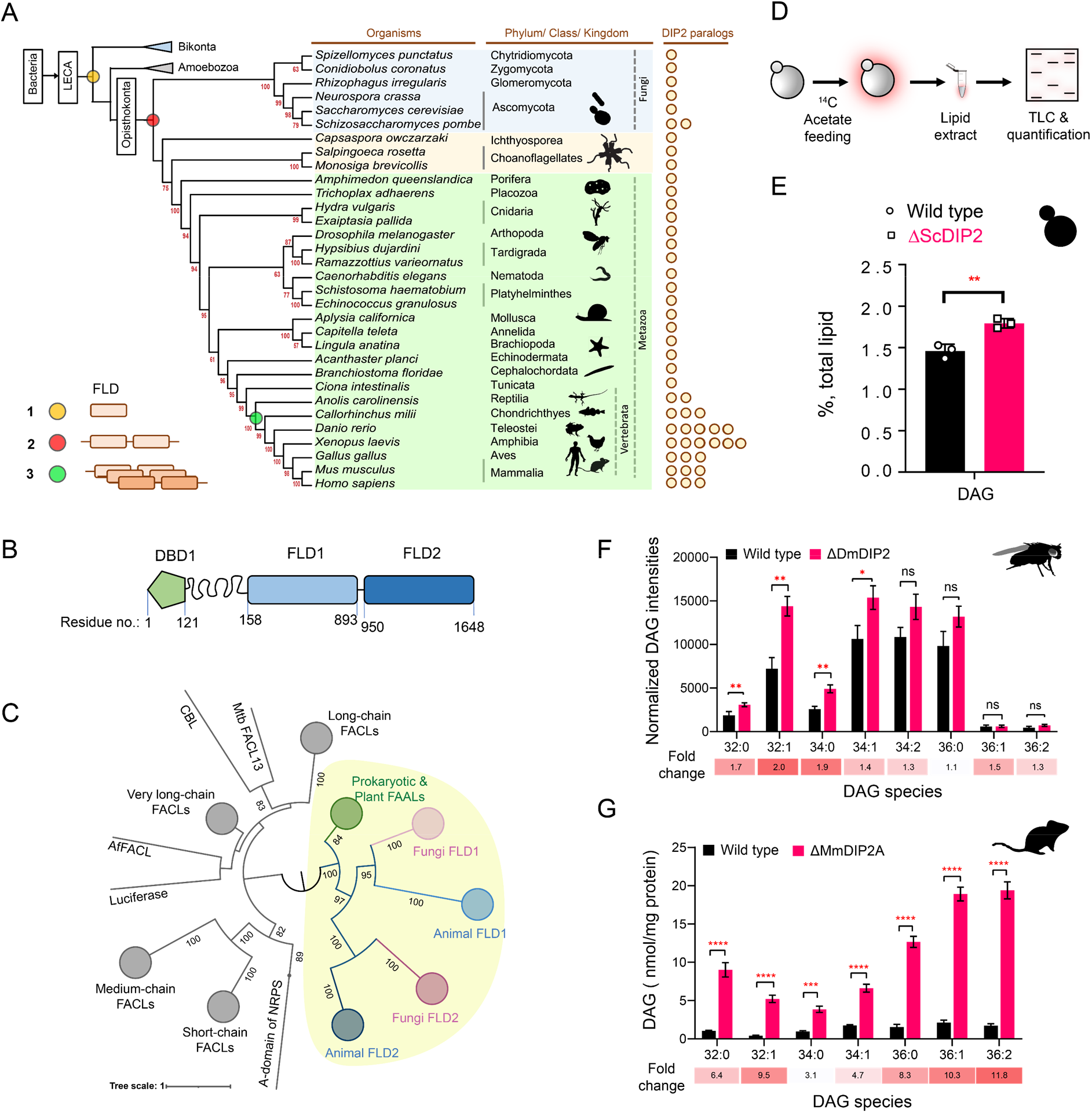
Tandem FAAL-like domains containing protein DIP2 is a new player in the DAG metabolism of Opisthokonta. (**A**) The distribution of FAAL-like domains in Eukaryota: three events in FLD evolution are indicated on a representative phylogenetic tree, (1) recruitment of bacteria-like standalone FLDs in eukaryotes (yellow circle), (2) domain duplication resulting in tandem FLDs in opisthokonts (red circle), and (3) whole gene duplications resulting in paralogs in vertebrates (green circle). (**B**) Domain architecture and predicted domain boundaries of yeast DIP2 homolog. Amino acid numbers are indicated for each domain boundary. DBD1-DMAP1 Binding Domain 1; FLD1-FAAL-like Domain 1; FLD2-FAAL-like Domain 2. (**C**) Phylogenetic distribution of ANL family of enzymes shows the close association of FLDs of DIP2 (FLD1 and FLD2) with prokaryotic/plant FAAL and divergence from FACLs of both prokaryotic and eukaryotic origins (nodes in grey colour). FAAL-like domain cluster is highlighted in a yellow background. (**D**) A schematic for the metabolic labelling of the total lipid pool by feeding the yeast with radioactive [1-^14^C] acetic acid. The percentage for each lipid is calculated as a percentage of the total ^14^C-radiolabel incorporated in the lipidome. (**E**) The estimated DAG levels show accumulation in a ΔScDIP2 mutant yeast in comparison to a wild type. Data are representative of three independent experiments, mean ± SD. **p < 0.01. (**F**) LC-MS based quantification of DAGs from total lipid extract from 5-day old adult *Drosophila melanogaster* shows an accumulation of DAGs. (**G**) The accumulation of DAGs is seen in the lipidome isolated from the embryonic stem cells generated from MmDIP2a^-/-^ mice in CD1 background. In (**F** and **G**), the average of estimated DAGs is shown on y-axis and the specific lipid length (combined number of carbons in two acyl chains) of DAGs is indicated on x-axis. Data are represented as mean ± SEM (n = 6; unpaired, two-tailed student’s t-test; *p < 0.05; **p <0.01; ***p < 0.001; ns= not significant). The fold change for each DAG species is indicated below the bar graphs.

To begin the investigation, we generated DIP2 knock-out yeast (hereafter referred to as ΔScDIP2) using homologous recombination (**Supplementary Fig. 2A and B**). We hypothesized a lipidomic change in the ΔScDIP2 cells based on: (i) the presence of FLDs in ScDIP2 (Patil et al., 2021), which are fatty acid-activating domains, (ii) in mice, the deletion of DIP2A results in abnormal fat depositions and growth rates depending on their dietary lipid compositions (Kinatukara et al., 2020), (iii) analysis of the gene ontology of its genetic interactors revealed that a majority of them participate in lipid metabolism (**Supplementary Fig. 2C**). To address this, we radiolabelled the yeast lipids in a steady-state using ^14^C-acetate and assessed how it fluxes through various cellular lipid pools using thin-layer chromatography (TLC) (Fig. 1D). ΔScDIP2 cells showed a significant ∼23% increase in the total DAG levels compared to the wild type (Fig. 1E) with a moderate depletion in TAG level (**Supplementary Fig. 2D**). However, none of the other membrane lipid classes showed any significant change (**Supplementary Fig. 2D**). This observation for the first time implicates the involvement of DIP2 in lipid metabolism, particularly DAG metabolism in yeast.

Subsequently, we probed other members of Opisthokonta to see if the identified role of DIP2 in DAG metabolism is conserved. We used mutants of DIP2 generated in other model organisms such as *Drosophila melanogaster* (ΔDmDIP2) (Nitta et al., 2017) and *Mus musculus* (ΔMmDIP2A) (Kinatukara et al., 2020). A comparative liquid chromatography–mass spectrometry (LC-MS) analysis of whole-tissue lipid extracts from adult wild type (Canton-S) and ΔDmDIP2 *Drosophila* showed ∼two-fold accumulation in a subset of DAG species (Fig. 1F). Interestingly, the absence of even a single paralog, DIP2A in the ES cells isolated from ΔMmDIP2A mice, showed ∼3 to 12-fold accumulation of DAG species compared to wild type ES cells (Fig. 1G). We noted a preferential accumulation of C16 acyl chain-containing subset of DAGs (C32:0, C32:1, C34:0, C34:1) in ΔDmDIP2 flies, while the accumulation is biased towards C32:1, C36:1 and C36:2 DAGs in ΔMmDIP2A ES cells (fold change indicated in Fig. 1F and G). Taken together, these data underscore the conserved role of DIP2 in regulating the levels of DAG from fungi to mammals.

### DIP2 regulates chemically distinct DAG subspecies in yeast by facilitating their conversion to TAG

To gain further insight into the function of DIP2 in DAG metabolism, we focused on yeast as the lipid metabolism pathways are well-documented. LC-MS analysis of yeast lipidome revealed a striking ∼19-fold accumulation of C36:0 and ∼8-fold accumulation of C36:1 chain-length DAG subspecies in ΔScDIP2 strain (Fig. 2A **and Supplementary Fig. 3A**). Genetic complementation with ScDIP2 under native or galactose-inducible (with a C-terminal GFP tag) promoters restored the levels of DAGs (Fig. 2B). To our surprise, the abundant DAG species (representing > 90% of total DAG pool), comprised of C32 and C34 chain-lengths (Casanovas et al., 2015; Ejsing et al., 2009), hereafter referred to as bulk DAG pool, remained unchanged in ΔScDIP2. Interestingly, this indicates that the DAG subspecies accumulation is independent of the well-characterized bulk DAG metabolizing pathways through Dga1, Lro1, Are1, Are2 (DAG acyltransferases) and Dgk1 (DAG kinase) (Li et al., 2020; Rockenfeller et al., 2018). Therefore, we conclude that DIP2 is required for the regulation of a defined subset of the DAG pool in yeast.

**Fig. 2.**
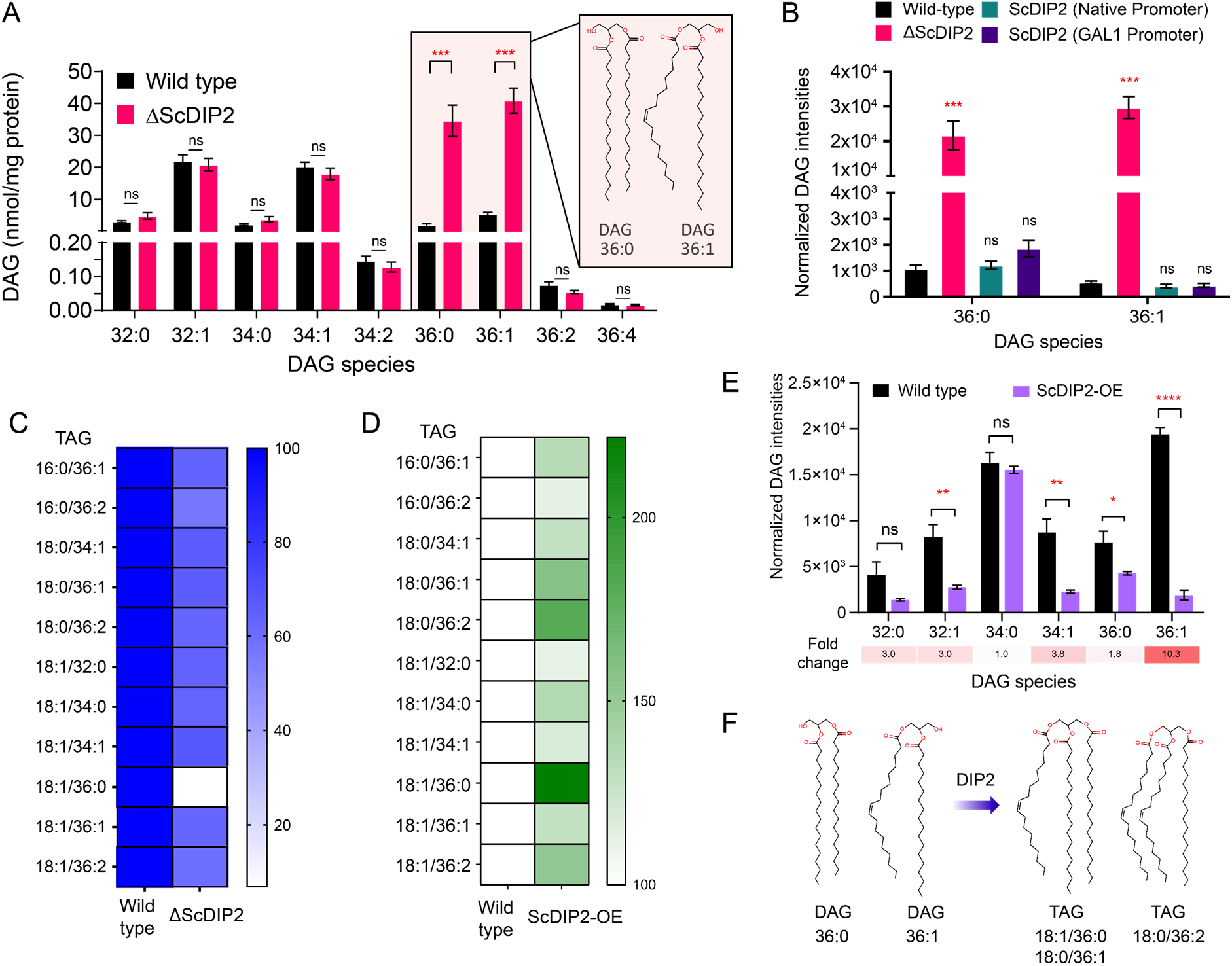
DIP2 regulates selective DAG pools by facilitating their conversion to corresponding TAGs. (**A**) LC-MS analysis of the ΔScDIP2 reveals a massive accumulation of C36:0 and C36:1 DAGs. Chemical structures of indicated DAGs are shown in the inset. (**B**) Genetic complementation of the mutant with native and galactose-inducible promoter-driven ScDIP2 expression. (**C**) Deletion of ScDIP2 resulted in depletion of TAGs. The decreased level of TAGs with various chain-lengths in ΔScDIP2 are shown as a linear gradient (n=6) and normalized to the corresponding wild type level (set at 100%). (**D**) Overexpression of ScDIP2 (GAL1 promoter) leads to accumulation of TAG species. The increased level of TAGs with various chain-lengths in ScDIP2-OE are shown as a linear gradient (n=6) and normalized to the corresponding wild type level (set at 100%). (**E**) The DAG levels reduced significantly below the basal level upon overexpression of ScDIP2. All data are represented as mean ± SEM (n>5; unpaired, two-tailed student’s t-test; *p < 0.05; **p < 0.01; ***p < 0.001; ****p < 0.0001; ns= not significant). The fold change for each DAG species is indicated below the bar graphs. (**F**) An illustration summarizing the role of DIP2 in facilitating the conversion of chemically distinct DAG to TAG (depicted by respective chemical structures).

DAGs are the central lipid intermediates that are channelled through *de novo* and salvage pathways to form diverse membrane and storage lipids (**Supplementary Fig. 3B**) (Carrasco & Merida, 2007). So, there is a possibility of C36:0 or C36:1 subset enrichment in other lipid classes of the ΔScDIP2 strain. However, the lipidomics profile of major phospholipids showed no alteration either in a chain-length dependent or independent manner (**Supplementary Fig. 3C-H**). Interestingly, metabolic radiolabelling showed a moderate but significant depletion (∼29%) of TAG level in ΔScDIP2 (**Supplementary Fig. 2D**), which is a storage lipid and a major reservoir for cellular DAG pools. Lipidomics revealed a depletion of ∼30-40% TAGs related to a defined subset of DAGs (C36:0 and C36:1), with the highest depletion of ∼93% in the case of C18:0/36:1 TAG (Fig. 2C **and Supplementary Fig. 4A**). The depletion could result either from an inefficient acylation of DAGs to TAGs or increased lipolysis of those TAGs to DAGs. The latter was ruled out as the LC-MS based quantification of the downstream products of TAG lipolysis (i.e., monoacylglycerols, MAG and free fatty acids, FFA) remained unchanged (**Supplementary Fig. 4B and C**). Taken together, the results suggest that ScDIP2 is involved in selective DAG to TAG flux, possibly by direct or indirect facilitation of an acylation reaction of DAG subspecies (**Supplementary Fig. 4D**). The depletion of TAG levels in ΔMmDIP2A mouse ES cell lines is in consensus with this argument (**Supplementary Fig. 5A**). However, it should be noted that the TAG levels of ΔDmDIP2 *Drosophila* remained comparable to wild type (**Supplementary Fig. 5B**), which perhaps could be attributed to a masking effect from the huge amount of TAG sourced from the fat-body of *Drosophila* (Heier & Kuhnlein, 2018).

We asked whether DIP2 has a direct effect on TAG synthesis from selective DAG subspecies. To test this, we overexpressed ScDIP2 (under GAL1 promoter), hereafter referred to as ScDIP2-OE, and checked the level of DAGs and TAGs. We found that the TAGs related to selective DAGs, C18:1/36:0 TAG increased the most (∼ 122%), while C18:0/36:1 and C18:0/36:2 TAG increased by ∼ 58 % and ∼ 80%, respectively in ScDIP2-OE cells (Fig. 2D **and Supplementary Fig. 4A**). Concomitantly, selective DAG species, C36:1 DAG showed the highest fold change (∼10-fold) in depletion, further below the basal level in ScDIP2-OE cells (Fig. 2E). It should be noted here that a few other DAG subspecies were also depleted significantly but less drastically than the selective DAG. Such promiscuity could be a result of non-physiological levels of ScDIP2 due to overexpression. Therefore, the data suggest that the DIP2 expression is crucial to maintain the basal level of DAG subspecies (Fig. 2F) and directly facilitates a DAG to TAG conversion process.

### DIP2-mediated selective DAG regulation protects yeast from ER stress

We then sought to decipher the physiological relevance of DAG subspecies regulation by DIP2 in yeast. As ΔScDIP2 cells showed normal growth and no distinct morphological defects under standard nutrient conditions (**Supplementary Fig. 6A and B**), we assessed its adaptability to different stress conditions such as nutrient deficiency, osmotic stress, temperature changes, redox stress to name a few (**Supplementary Fig. 6C**). ΔScDIP2 cells showed significant growth impairment in the presence of tunicamycin (Fig. 3A), which was earlier observed in a genome-wide screen (Chen et al., 2005). Tunicamycin, an inhibitor of *N*-linked glycosylation in the endoplasmic reticulum (ER), induces misfolded protein accumulation in ER lumen leading to proteostasis defect, commonly referred to as ‘ER stress’. Genetic complementation using native promoter (Fig. 3A) or GAL1 promoter (**Supplementary Fig. 7A and B**) was sufficient to rescue tunicamycin-induced ER stress. ER stress is known to activate UPR signalling, a highly conserved homeostasis signalling in all eukaryotes (Han et al., 2010; Ron & Walter, 2007). Therefore, we assessed the levels of UPR activation, using fluorescence-activated cell sorting (FACS)-based assay (Jonikas et al., 2009; Merksamer et al., 2008; Surma et al., 2013). The intensity of the fluorescence signal from the reporter is a direct measure of the levels of UPR activation when the cells are subjected to ER stress and the assay was validated by titrating with tunicamycin (**Supplementary Fig. 6D and E**). ΔScDIP2 cells exhibited a ∼1.6-fold elevated UPR stress than basal levels found in wild type (Fig. 3B) and the addition of tunicamycin only exacerbated the already active ER stress in the mutant **(Supplementary Fig. 6F**). It implies that the mutant suffers from ER stress even in the absence of stress-inducing agents.

**Fig. 3.**
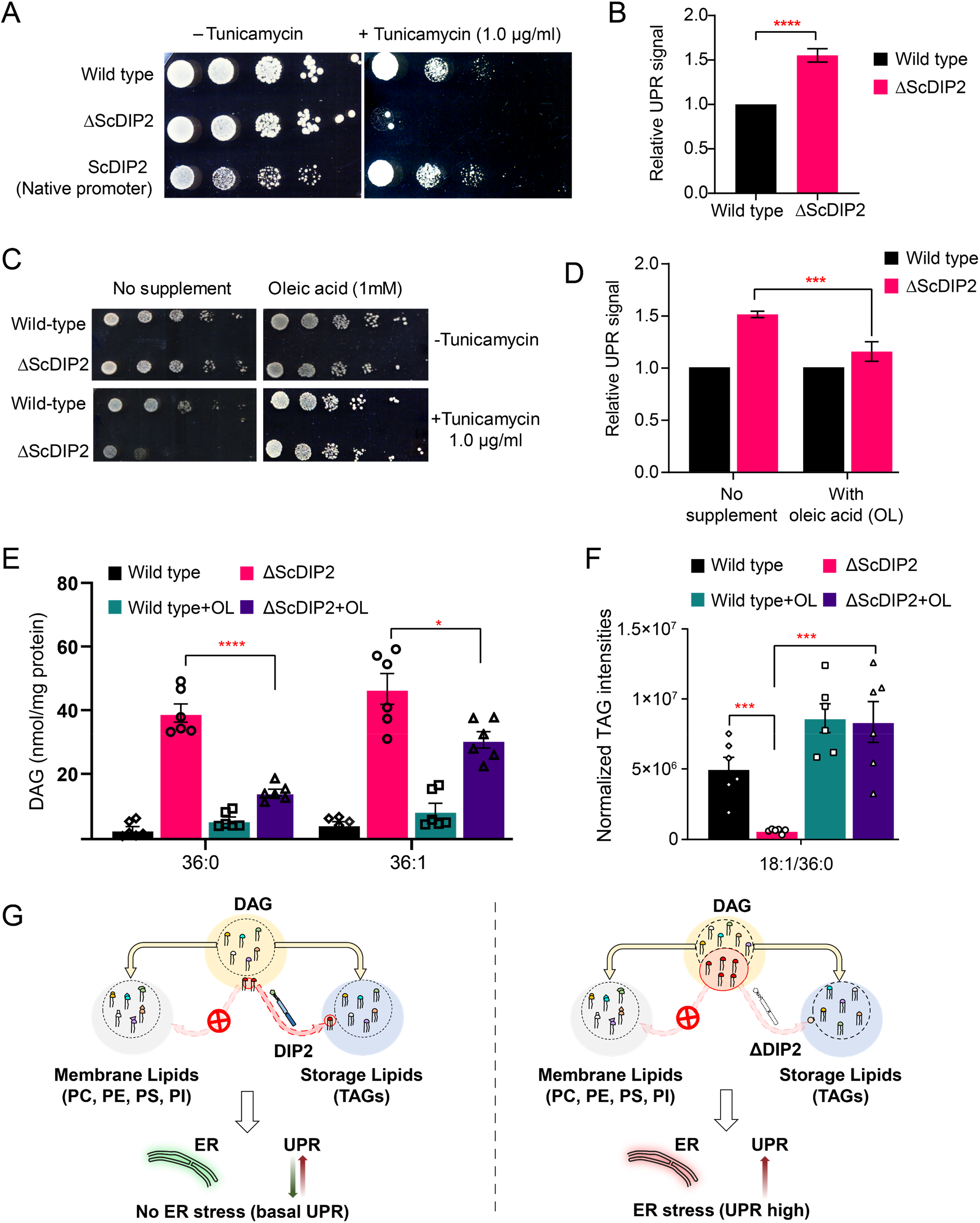
Restoration of selective DAG level alleviates constitutive ER stress in DIP2 mutant yeast. (**A**) Serial dilution assay of yeast with indicated strains show that ΔScDIP2 cells are sensitive to Tunicamycin-induced ER stress. A representative image of the assay where expression of ScDIP2 using native promotor (genetic complementation) rescuing ER stress in ΔScDIP2 is shown. (**B**) The 4X-UPRE reporter-based quantification of UPR signalling level. The fractional increase in RFP signal intensity (UPR signal) measured in the mutant cells from the log phase relative to wild type cells in the log phase is represented. Data are represented as mean ± SD (unpaired, two-tailed student’s t-test; n = 6; ****p > 0.0001). (**C**) Serial dilution assay with chemical supplementation of OL (1 mM) shows rescue of ER stress sensitivity in ΔScDIP2 cells. (**D**) 4X-UPRE reporter assay wild type and ΔScDIP2 grown with 1 mM OL. The data shows a reduction in UPR signal in ΔScDIP2 cells when supplemented with OL. Data are represented as mean ± SD (n = 4; unpaired, two-tailed student’s t-test; ***p < 0.001). (**E** and **F**) The lipidomics analysis of the oleic acid supplemented ΔScDIP2 cells shows that the accumulated C36:0 and C36:1 DAG species are significantly reduced. Corresponding TAG species (C18:1/36:0) is also normalized to basal level upon oleic acid supplementation. Data are represented as mean ± SEM (n > 5; unpaired, two-tailed student’s t-test; *p < 0.05; **p < 0.01; ***p < 0.001; ****p < 0.0001; ns= not significant). (**G**) A schematic representing the redirection of the selective DAG pool (red circle) to TAG by ScDIP2 helps in balancing the compositional diversity of DAG which is critical for ER homeostasis. Bulk DAG pool is utilized for membrane lipid and storage lipid (TAG) generation (shown via yellow arrow) while selective DAG pool can only be redirected to TAG (shown via red arrow).

Since the link between UPR and lipid metabolism is well-established (Volmer & Ron, 2015; Volmer et al., 2013), aberrant DAG metabolism in ΔScDIP2 cells could be ascribed for the ER proteostasis defect. Therefore, we hypothesized that by metabolically fluxing these DAGs into other lipids, we may simply be able to alleviate the ER stress in ΔScDIP2 cells. It is known that specific lipid precursors such as choline, ethanolamine and FAs can facilitate the utilization of cellular DAG pool to produce phosphatidylcholine (PC), phosphatidylethanolamine (PE), and TAGs, respectively (Boumann et al., 2004; Henneberry et al., 2001; Montell et al., 2001). Therefore, we performed a “chemical complementation” screening to rescue the ER stress in ΔScDIP2 cells by supplementing with lipid precursors. We found that except for oleic acid (OL; C18:1 FFA), none of the other precursors rescues the ER stress phenotype (Fig. 3C **and Supplementary Fig. 7C**). The supplementation of ΔScDIP2 mutants with a different unsaturated FA (e.g.: palmitoleic acid; C16:1 FFA) or saturated FA (e.g.: palmitic acid; C16:0 FFA, stearic acid; C18:0 FFA) did not relieve ER stress (**Supplementary Fig. 7D**). Furthermore, OL supplementation is enough to reduce the constitutive UPR upregulation in ΔScDIP2 (Fig. 3D). In agreement with those observations, the LC-MS analysis revealed the restoration of the accumulated subset of DAGs back to basal levels (Fig. 3E). All the other precursors showed little or no effect on the levels of an accumulated subset of DAGs (**Supplementary Fig. 7E**), which also correlated with their inability to suppress tunicamycin sensitivity. A concomitant normalization in related TAGs was also observed upon OL supplementation only (Fig. 3F **and Supplementary Fig. 7F**). It is interesting to note that bulk DAG accumulation is possibly less toxic to ER as the null mutants of canonical DAG-utilizing enzymes such as Dga1 and Lro1 or Dgk1 are not sensitive to ER stress (**Supplementary Fig. 8**). This indicates the atypical role of selective DAG subspecies as their accumulation results in ER stress, while ScDIP2 alleviates the ER stress by facilitating the buffering of such DAGs from the membrane pool to the storage pool (Fig. 3G).

Since OL supplementation was found to restore DAG level, we wondered if any physiological condition(s) can mimic this process. Under nutrient limitation conditions such as stationary phase, yeast is known to massively upregulates endogenous OL production and subsequent TAG synthesis, facilitated by majorly Dga1 and Lro1, as a metabolic strategy to store energy in the form of neutral lipids (Casanovas et al., 2015; Jacquier et al., 2011; Joshi et al., 2018; Markgraf et al., 2014). To test this, the DAG levels in the early-log, mid-log and late stationary phases of wild type were compared with the levels in ΔScDIP2 cells. The bulk DAG levels across the various stages of growth phases remain constant in the wild type, which is also observed in earlier reports (Casanovas et al., 2015). Strikingly, the ΔScDIP2 cells showed the accumulation of a selective subset of DAGs only during the early and mid-log phase and these levels are restored during the stationary phase (Fig. 4A). Thus, the toxic accumulation of selective DAGs is diluted by the bulk conversion of DAGs to TAGs activated during the stationary phase, which also suppress the activated UPR (Fig. 4B). It is, therefore, evident that the role of ScDIP2 in regulating a defined subset of DAGs to TAG flux must be a growth phase dependent phenomenon. So, we tracked the expression of ScDIP2 along the growth phase using a ScDIP2-GFP knock-in line (C-terminal GFP). The expression of ScDIP2-GFP appeared as an intra-cellular tubular and punctate-like structure (**Supplementary Fig. 9A**) only during the early-log to mid-log phase and rapidly diminished in the stationary phase (Fig. 4C). Taken together, these observations reveal an interesting parallel between canonical and ScDIP2-dependent regulation of DAG subspecies required to protect ER function in the logarithmic growth phase (Fig. 4D).

**Fig. 4.**
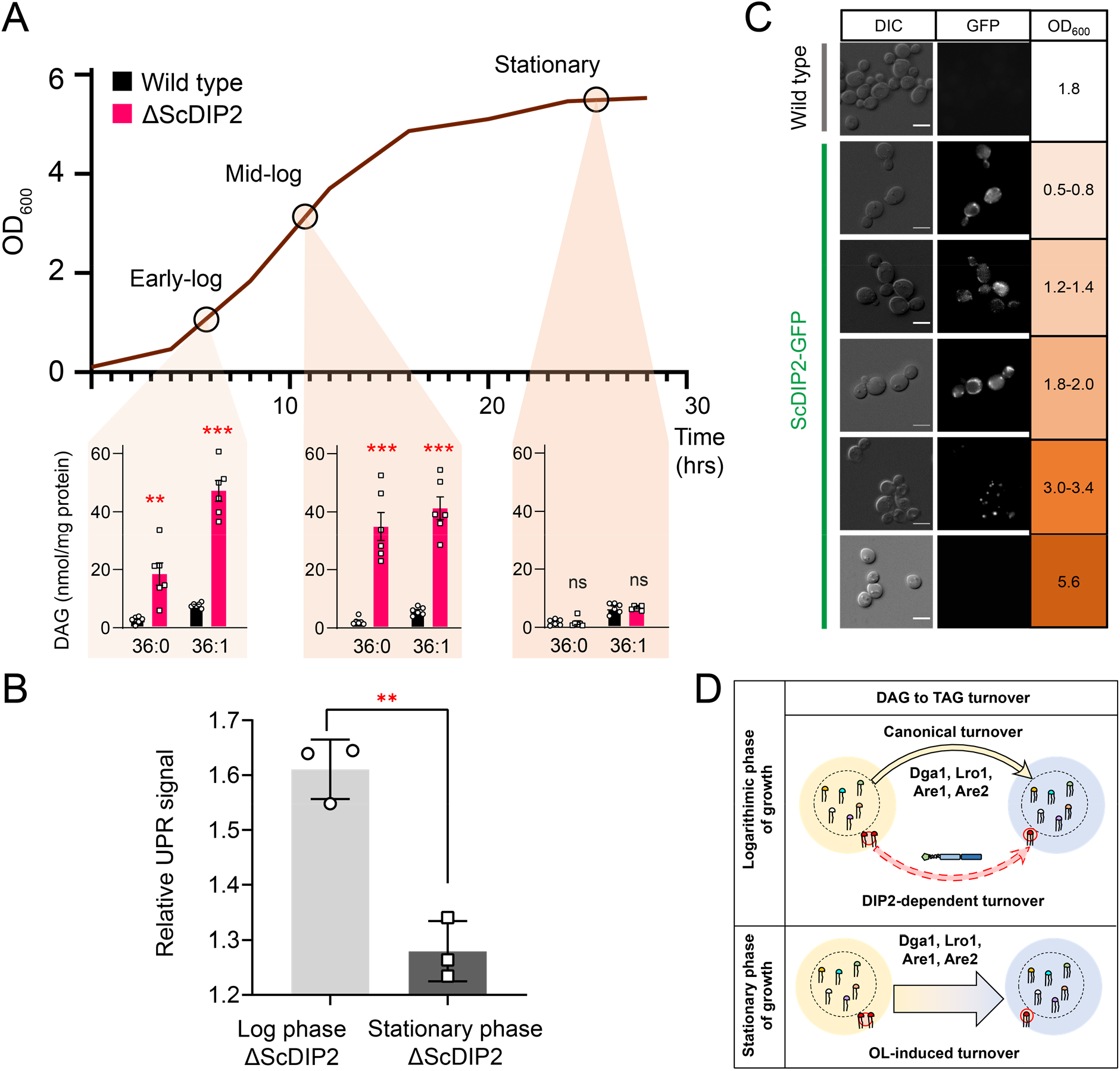
Growth phase-dependent regulation of selective DAGs by DIP2 ensures ER homeostasis in yeast. (**A**) Tracking the level of selective DAGs in ΔScDIP2 cells across the growth phases: early-log, mid-log, and stationary phase. Data are represented as mean ± SEM (n > 5; unpaired, two-tailed student’s t-test; **p < 0.01; ***p < 0.001; ****p < 0.0001; ns= not significant). (**B**) The 4X-UPRE reporter assay shows a reduced level of UPR activation when ΔScDIP2 cells are grown till the stationary phase. Data represents fold change in ΔScDIP2 relative to respective wild type control (mean ± SD; n=3; unpaired, two-tailed student’s t-test; **p < 0.01). (**C**) Growth-phase dependent expression of ScDIP2 is shown via the fluorescence signal of GFP tagged with C-terminal ScDIP2 in its genomic locus. Scale bar = 5 µm. (**D**) A schematic showing DIP2-dependent selective DAG to TAG turnover is operative in the log phase while a surge in OL-induced TAG biosynthesis during the stationary phase restores the overall DAG level.

### DIP2-regulated selective DAG species are required for vacuole-fusion mediated osmoadaptation

Since DAGs are metabolized at multiple organellar sites (Baron & Malhotra, 2002; Cowell et al., 2009; Ganesan et al., 2019; Starr & Fratti, 2019; Yang & Kazanietz, 2003), the sub-cellular location of ScDIP2 may indicate the physiological niche of selective DAGs. The ScDIP2-GFP showed co-localization with FM4-64, a vacuolar marker, as distinct patches on the vacuolar surface (Fig. 5A and B). In addition, the ScDIP2-GFP signal was also found to be associated with a mitochondrial marker, MitoTracker^TM^ (**Supplementary Fig. 9C-H**). A crude cell fractionation also showed the association of ScDIP2 with the total membrane fraction isolated from log-phase cells (**Supplementary Fig. 9B**). Given the localization of ScDIP2 to vacuoles and its role in DAG regulation, we hypothesized that loss of ScDIP2 would affect vacuole function. Therefore, we examined the vacuole morphology in the log phase as it serves as an index of vacuole health. We found that about 60% of ΔScDIP2 cells contain a greater number of single, rounded and enlarged vacuoles, which is roughly twice that of the wild type (Fig. 5C and D). A comparative analysis of the size distribution of vacuoles in the log phase of growth showed that larger sized vacuoles are more frequently observed in ΔScDIP2 mutant than the wild type (Fig. 5E). Thus, the increase of single and large vacuoles in ΔScDIP2 cells is presumably due to aberrant vacuole membrane fusion driven by accumulated DAGs, a known membrane fusogen (Starr & Fratti, 2019). The optimum vacuole fusion is critical for cells’ adaptation to osmotic stress, a common environmental stress experience by fungi (Veses et al., 2008), To check the functional consequence of such compromised vacuolar homeostasis, ΔScDIP2 cells were subjected to hyper-osmotic stress using high-salt growth media. An incubation in hyper-osmotic conditions resulted in two-to three-fold higher cell death in ΔScDIP2, while the cell viability was identical for both the strains in the absence of stress (Fig. 5F). A similar observation was also made during the screening of ΔScDIP2 using serial dilution assay (**Supplementary Fig. 6C**).

**Fig. 5.**
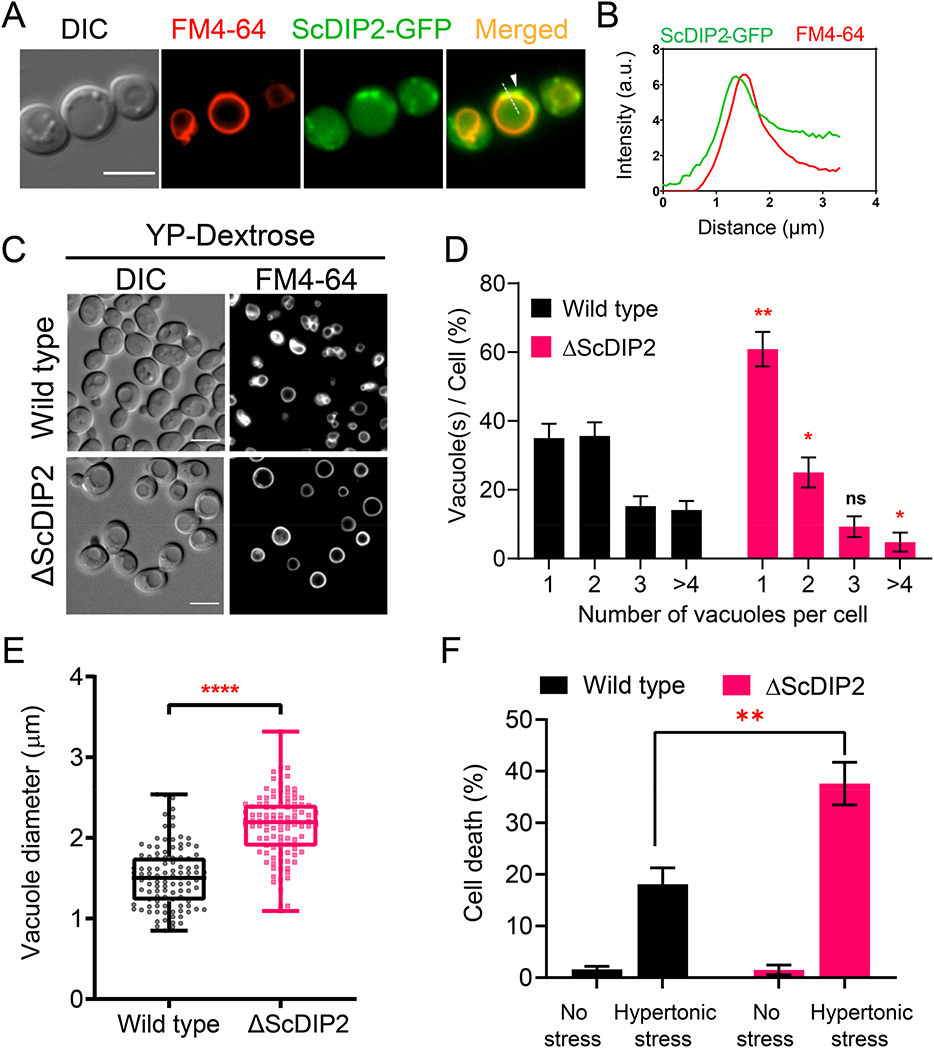
Loss of DIP2 results in aberrant vacuolar fusion leading to failure in osmoadaptation. (**A**) Live cell imaging of ScDIP2-GFP knock-in strain for subcellular localization study. ScDIP2-GFP cells (mid-log phase) are stained with FM4-64 (red) dye and images have been obtained using respective channels in a wide-field epifluorescence microscope. White arrowhead denotes co-localization site. Scale bar = 5 µm. (**B**) Line scan of signal intensity along the white line (near white arrowhead) shown in (A). (**C**) Representative images of vacuole morphology of the indicated strains grown till log phase in YP-dextrose media. Cells were stained with FM4-64. Scale bar = 5 µm. (**D**) Quantitation of vacuoles observed per cell of indicated strains imaged in (C). Data were represented as mean ± SD (n = 3; > 100 cells per strain; unpaired, two-tailed student’s t-test; *p < 0.05; **p < 0.01; ns= not significant). (**E**) Vacuole diameter has been measured by line-scan analysis of FM4-64-stained vacuoles from yeast cells at the budding stage. Each dot represents a single vacuole per cell (>100 cells per strain) and box and whisker plot representing diameters (in µm) of the largest vacuole of indicated strains. Data are representative of at least three independent experiments (unpaired, two-tailed student’s t-test; ****p<0.0001). (**F**) ΔScDIP2 and wild type cells are subjected to hypertonic stress and then stained with trypan blue for assessing cell viability. Data are represented as mean ± SD (n > 3; unpaired, two-tailed student’s t-test; *p < 0.05; **p < 0.01; ****p < 0.0001; ns= not significant).

Next, we investigated if DAG depletion upon ScDIP2 overexpression can show converse effects on vacuole morphology. The vacuoles in the ScDIP2-OE strain were found to be highly fragmented; single vacuole containing cells are about three times lower while the multi-vacuolar (> 4) cells were three times more than that in the wild type (Fig. 6A and B). To confirm whether this fragmented vacuolar morphology is due to the impairment of vacuole fusion, we employed a vacuole fusion-fission assay utilizing the osmotic response of yeast (Fig. 6C). The growth in hypertonic media-induced vacuole fission in yeast, resulting in highly fragmented vacuoles in both wild type and ScDIP2-OE cells (Fig. 6D and E). Exposure of such cells with fragmented vacuoles to hypotonic conditions induces rapid vacuole fusion. Vacuoles in ScDIP2-OE cells are found to be highly fragmented and multi-lobar even after the fusion induction using hypotonic media (Fig. 6F), suggesting a significantly reduced vacuolar fusion process (∼ 30% reduced fusion compared to wild type) (Fig. 6G). Since rapid vacuole fusion is required during hypo-osmotic stress adaptation by yeast cells, we hypothesized that ScDIP2-OE cells will show reduced fitness in hypotonic conditions. Indeed, we found that acute hypotonic stress caused severe cell death in ScDIP2-OE while ΔScDIP2 cells showed significant tolerance to the stress compared to the wildtype (Fig. 6H). Together, the data suggest that the selective DAG pool is essential for the vacuole fusion process and ScDIP2-mediated regulation optimizes the process to ensure osmoadaptation in yeast.

**Fig. 6.**
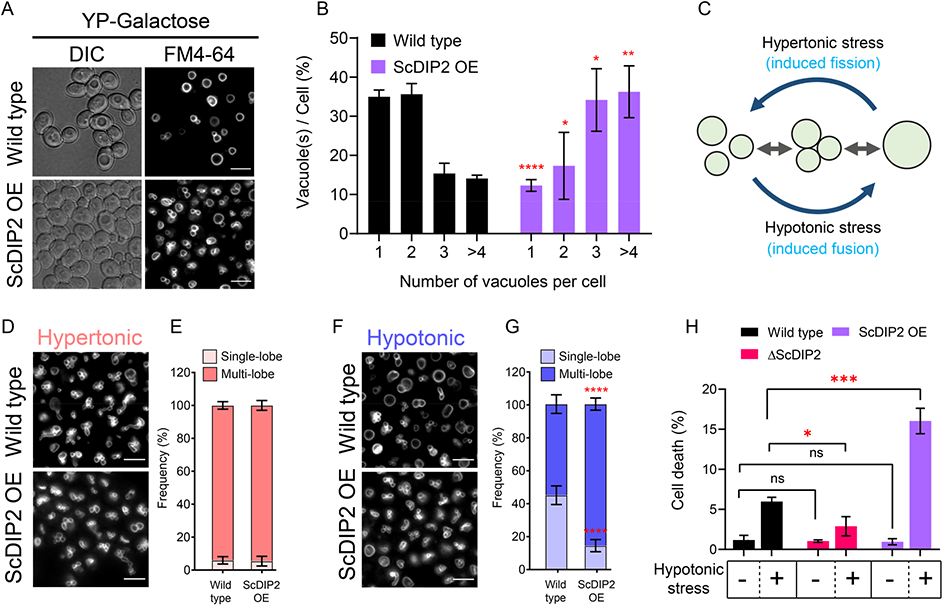
Depletion of DAG upon DIP2 overexpression destabilizes vacuolar fusion-fission homeostasis in yeast. (**A**) Representative images of vacuole morphology in indicated strains grown till log phase in YP-galactose media. Vacuoles are stained with FM4-64. Scale bar = 5 µm. (**B**) Quantitation of vacuoles observed per cell of indicated strains imaged in (a). Data are represented as mean ± SD (n = 3; > 100 cells per strain; unpaired, two-tailed student’s t-test; *p < 0.05; **p < 0.01; ****p<0.0001; ns= not significant). (**C**) A schematic depicting the vacuole fusion-fission assay utilizing the osmotic response of yeast to different concentrations of NaCl. (**D** and **F**) Representative images of FM4-64-stained vacuoles from indicated strains after treating with hypertonic (YPD + 0.4M NaCl) condition for 10 minutes, followed by hypotonic (water) condition for 10 minutes. (**E** and **G**) Quantification of vacuoles in hypertonic conditions (shades of pink) and hypotonic conditions (shades of blue). Frequency percentage of single-lobe (fused) and multi-lobe (fragmented) vacuole per cell from indicated strains imaged in D and F. Cells with 3 or more vacuoles were combined as ‘multi-lobe vacuole’ set (n = 6, > 100 cells per strain; unpaired, two-tailed student’s t-test; ****p < 0.0001). (**H**) Indicated strains are subjected to hypotonic shock by incubating in water for 10-15 minutes and then stained with trypan blue for cell viability. Data are represented as mean ± SD (n > 3: two-tailed student’s t-test; *p < 0.05; **p < 0.01; ****p < 0.0001; ns= not significant).

### Catalytic activity of tandem FLD-didomain is critical for DAG regulation function

Next, we asked whether the FLDs of DIP2 are required for the DAG regulation. Bioinformatics analysis on the emergence of the unique three-domain architecture of DIP2 in opisthokonts revealed that the N-terminal DMAP1-binding domain (DBD1) is absent in several ancient members of this group (viz. Nucleariids, Ichthyosporea, Choanoflagellates and some fungi species). However, the tandem FLD-didomain (FLD1-FLD2) architecture is conserved throughout all opisthokonts (Fig. 7A). For genetic complementation, we generated multiple GAL1-based domain-truncated constructs of ScDIP2 and their expression was validated by fluorescence microscopy and western blot experiments (**Supplementary Fig. 10A and B**). The genetic complementation assay with multiple domain-truncated constructs revealed that tandem FLD-didomain is minimally required to prevent ER stress, while the DBD1 is dispensable (Fig. 7B). We also performed lipidomics of ΔScDIP2 yeast complemented with different domain-truncated versions of ScDIP2. We found that the FLD-didomain restored the levels of the subset of DAGs characteristic to yeast (Fig. 7C), accompanied by the generation of related TAGs (Fig. 7D). Such a restoration of the DAG level is not evident when either of the FLDs is missing (**Supplementary Fig. 10C**). These results suggest that the recruitment and duplication of FLDs as an innovation in early Opisthokonta evolution to tackle a specific fraction of the DAGs pool, which if accumulated may become toxic.

**Fig. 7.**
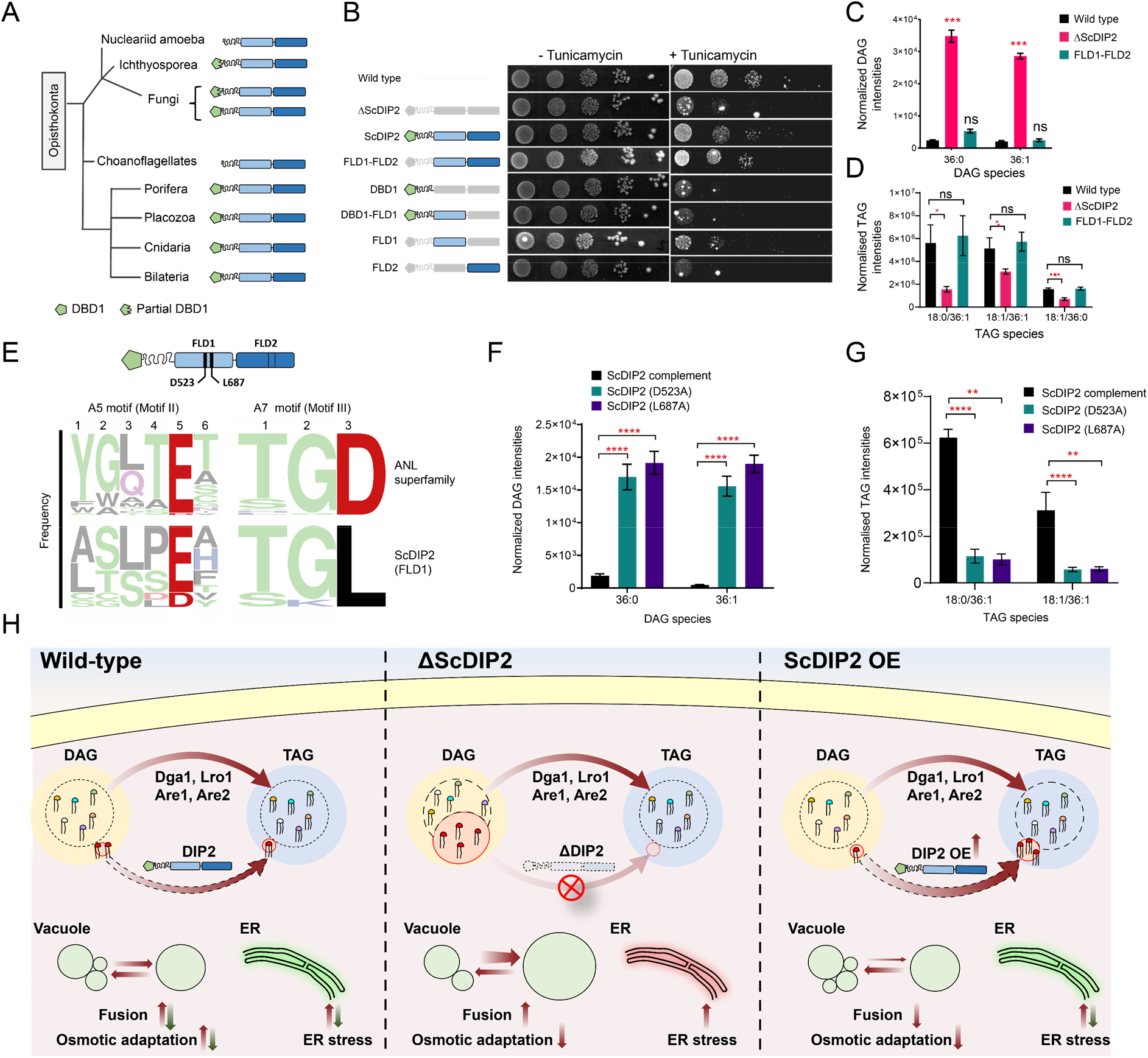
Catalytic activity along with tandem FLD-didomain organization is critical for the function of DIP2. (**A**) A graphical representation of the phylogenetic distribution of DIP2 across Opisthokonta showing conservation of FLD1-FLD2 didomain. (**B**) Tunicamycin-mediated ER stress rescue assay by complementing ΔScDIP2 with different domain-truncated versions of ScDIP2. Deleted domains are shown in grey colour. (**C and D**) Increased DAGs in ΔScDIP2 are restored to wild type level upon complementation with FLD1-FLD2 didomain, along with an increase in related TAG levels. (**E**) Graphical representation to show the crucial residues of adenylation motifs of ScDIP2 FLD1 and sequence logo images depicting the conservation of adenylation motifs of ANL superfamily in ScDIP2. (**F**) Adenylation mutants of ScDIP2 are not able to restore accumulated DAGs to basal level due to failure in fatty-acyl activation. (**G**) Selective TAG level is not restored to wildtype level upon expression on adenylation mutants of ScDIP2. Data are represented as mean ± SEM (n > 5; unpaired, two-tailed student’s t-test; *p < 0.05; **p < 0.01; ***p < 0.001; ****p < 0.0001; ns= not significant). (**H**) A model depicting the role of DIP2-mediated selective DAG regulation for cellular homeostasis and adaptation. The regulation of DAG subspecies, representing a small fraction (shown in red head-colours) of the total DAG pool without altering the levels of bulk DAGs (multiple head-colours) by redirecting them to TAGs (shown in red), is facilitated, by DIP2 in yeast. Such a regulation of minor DAG subspecies is essential for optimal vacuole (shown as green circle) fusion-fission dynamics leading to efficient osmotic adaptation and maintenance of ER homeostasis. In ΔScDIP2, the accumulation of these minor DAG subspecies results in ER stress, increased vacuole fusion and reduced osmotic adaptation capability. On the contrary, in ScDIP2-OE, these DAGs are depleted below the basal level, resulting in restored ER stress but reduced vacuole fusion along with the defect in osmotic adaptation.

To delineate if the FLDs have retained the enzymatic activity of a typical ANL superfamily module in DAG regulation, we performed sequence and mutational analysis of FLDs. A typical ANL superfamily member, including FLDs, should activate their respective substrates by the formation of an acyl-adenylate intermediate. All these homologs show highly conserved residues responsible for the acyl-adenylate formation, referred to as A5 and A7 motifs (Gulick, 2009). Structure-based sequence alignments, using models from AlphaFold Protein Structure Database (Jumper et al., 2021; Varadi et al., 2021), suggested that the A5 and A7 motifs are conserved in the FLD1 domains of all opisthokonts (Fig. 7E) and strikingly, FLD2 showed degeneration of both the motifs. This is also supported by the greater divergence of FLD2 domains from the prokaryotic/plant FAALs than FLD1 domains, as inferred from the branch-lengths of FLD clades in phylogenetic tree (Fig. 1C). Mutations in catalytic residues of adenylation motifs of FLD1 (Fig. 7E) led to the loss of function of ScDIP2. These adenylation mutants failed to rescue the ΔScDIP2 from tunicamycin-induced ER stress (**Supplementary Fig. 11D**) and also showed an inability to regulate the levels of accumulated DAGs by converting them to related TAGs (Fig. 7F and G). The expression and the localization pattern of FLD1 adenylation mutants were identical to the wildtype ScDIP2 (**Supplementary Fig. 11B and C**). The mutation of predicted residues of the degenerate motifs in FLD2 did not lead to loss of function (**Supplementary Fig. 11E**). Taken together, the adenylation activity of FLD1 of DIP2 is required for facilitating the conversion of selective DAGs to TAGs. Also, it can be suggested that FLD2 has not retained adenylation activity and has diverged for a yet to be identified function.

## Discussion

‘Why do cells possess a highly diverse lipidome?’ is one of the most perplexing questions in biology. Diversity amongst various classes of lipids arising from the variability in FA compositions such as chain-length, degree of unsaturation, is turning out to be crucial for biological functions (Atilla-Gokcumen et al., 2014; Khandelwal et al., 2021; Raghu, 2020; Shin et al., 2020). Earlier *in vivo* studies have pointed to the exclusive role of selective DAG subspecies in determining the fate of different physiological processes including signal transduction, cell polarity, membrane trafficking, ligand-binding to membrane receptors and modulation of membrane properties (Lee et al., 1991; Marignani et al., 1996; Schuhmacher et al., 2020; Ware et al., 2020). However, the molecular players and pathways in selective regulation are largely unknown. Interestingly, studies in mammals have revealed the presence of a unique isoform of diacylglycerol kinase, namely DGKε, that channelizes specific DAG species, suggesting the evolution of selective metabolism routes in eukaryotes (Milne et al., 2008). However, the known DAG metabolizing enzymes of yeast, DAG acyltransferase (Dga1, Lro1, Are1 and Are2) and DAG kinases (Dgk1), are non-specific in nature and thus facilitate a generic regulation through bulk conversion of DAGs (Li et al., 2020; Rockenfeller et al., 2018). Therefore, a thorough investigation is needed to identify how ∼2 billion years of eukaryotic evolution has shaped the lipid repertoire through selective regulation mechanisms. In this context, our study demonstrates the cellular function of a conserved protein family, DIP2 in regulating a subset of DAG pool (C36:0 and C36:1 DAGs) of yeast in a growth phase dependent manner. In yeast, the loss of DIP2 leads to the accumulation of these DAG subspecies and a concomitant depletion of chain-length related TAG species. These DAG subspecies are found to be toxic to yeast cells as their accumulation in the absence of DIP2 interfere with cellular processes like ER homeostasis and osmoadaptation. Overexpression of DIP2 in yeast increases the flux from DAG to TAG, resulting in massive depletion of DAG subspecies below the basal level. As we find that these DAG subspecies are required for vacuole membrane fusion, diminishing their cellular abundance upon DIP2 overexpression leads to decreased vacuole fusion and increased cell death upon osmotic stress in yeast. Thus, the study essentially identifies an unconventional and selective DAG metabolism route in yeast with important physiological consequences. Similar to the yeast results, the lipidomics analysis indicates that the DIP2-mediated DAG regulation in *Drosophila* and mice is also biased towards a subset of DAG species. Given the tissue-specific expression (Nitta et al., 2017; Zhang et al., 2015) and the presence of multiple paralogs of DIP2 in higher organisms, the specificity and the functional significance of DAG regulation by DIP2 need to be probed in further studies.

The study characterizes DIP2 as a noncanonical member of DAG metabolism pathways as it shares no homology with known DAG metabolizing enzymes of eukaryotes. Interestingly, DIP2 harbours a tandem FLD-didomain which itself is a novel domain organization with respect to the evolutionary history of FLD. Although the canonical function of standalone FLD is to activate fatty acids, we find that the didomain architecture of FLD is essential for the DAG to TAG conversion by DIP2. We also provide evidence for the role of adenylation activity of FLDs of ScDIP2 in facilitating the conversion of DAG to TAG using mutations in the conserved adenylation motifs. This leads us to propose a putative biochemical scheme where DIP2 is involved in the fatty acylation of DAGs to produce TAGs, utilizing an activated acyl-adenylates intermediate. Despite significant efforts, we were unable to show the DAG to TAG conversion reaction *in vitro* using individual components along with purified protein. Although this marks the limitation of the current study, it highlights the involvement of other cellular factors or specific membrane environment necessary for the reaction. The involvement of known DAG acyltransferases (Dga1, Lro1, Are1 and Are2) in the DIP2-mediated DAG regulation pathway cannot be ruled out. However, the accumulation of the subset of DAGs in a DIP2 mutant despite the presence of those acyltransferases does suggest that DIP2 mediated pathway is independent of known TAG forming enzymes (Fig. 7H). Since DAG to TAG conversion via canonical pathways takes place at ER and lipid droplets (Sorger & Daum, 2003), the association of ScDIP2 with vacuole and mitochondria begs further investigation into the possibility of the noncanonical nature of DIP2-mediated DAG metabolism pathway.

Total DAG pool constitutes only a relatively small population (3-5%) within the lipidome (Casanovas et al., 2015; Ejsing et al., 2009) and is maintained via the rapid turnover to either storage lipids or membrane lipids. Indeed, the deletion of DAG metabolizing enzymes of yeast such as Dgk1, which facilitates the formation of phospholipids, or Dga1 and Lro1, that facilitate TAG synthesis results in the accumulation of bulk DAG species (Fakas et al., 2011; Mora et al., 2012; Oelkers et al., 2002; Oelkers et al., 2000; Rockenfeller et al., 2018). Surprisingly, unlike the DIP2 mutant which accumulates a subset of the DAG pool (< 0.005%), the accumulation of bulk DAGs in Dga1 or Lro1 or Dgk1 mutants does not lead to ER stress induction. It evidently points towards functionally independent DAG populations, viz., a benign bulk DAG pool, utilized mainly for bilayer production and energy storage, and a minor context-dependent DAG pool. The present study identifies the minor DAG pool as being resistant to any kind of redirection that retains them in the membrane. These DAGs can only be titrated as a storage lipid, TAG, by DIP2 in the logarithmic growth phase and via oleic acid-mediated ‘metabolic reset’ in the stationary phase. Thus, the accumulation of these DAG subspecies is likely to impact the physical properties of the membrane, which subsequently results in UPR triggering and vacuole fusion dysregulation.

Overall, the current work shows that DIP2-mediated fine-tuning of a fraction of the DAG pool is required for adapting to different types of stress such as ER proteostasis and osmolarity. Unlike the laboratory conditions, these adaptation helps in the survival of yeast in the diverse environmental niches in nature (Fig. 7H). As pathogenic fungi adapt to the changing microenvironments during disease progression, similar stress adaptation processes linked to DAG regulation becomes crucial for their survival inside the host. Studies on plant and animal pathogenic fungi have indeed suggested that DIP2 is involved in infection-related morphogenesis like conidia, appressoria, hyphae and spherule formation (Lu et al., 2003; Narra et al., 2016; Wang et al., 2016), where vacuole fusion-fission dynamics plays a vital role (Veses et al., 2008). Consequently, DIP2 has been identified as a potential candidate for designing fungicides and developing a vaccine against valley fever (Shubitz et al., 2018). Furthermore, DAG regulation in neurons is found as a critical factor for neuron branching, dendritic spine formation (Tu-Sekine et al., 2015), etc., which are the major cellular processes affected in autism and related neurodevelopmental disorders (Gilbert & Man, 2017; Phillips & Pozzo-Miller, 2015). Since DIP2 has been recognized as a potential risk factor for ASD and other neurodevelopmental disorders (**Supplementary Table 1**), aberrant accumulation of chemically distinct DAG subspecies is likely to be the potential molecular driver of these disorders. Interestingly, the previous reports suggesting the requirement FLD-didomain of DIP2 for proper neuron branching in *Drosophila* and *C. elegans* (Nitta et al., 2017; Noblett et al., 2019) correlates with our observation on the need of FLD-didomain in selective DAG to TAG conversion. This also suggests the possible involvement of DIP2 in driving these crucial developmental processes via DAG subspecies regulation. Therefore, the identification of DIP2 as a regulator of DAG subspecies abundance provides a solid platform to bridge the gap between its cellular function and phenotypes linked to disease pathogenesis. Further explorations will be focussed on, (i) the elucidation of the biochemical and structural mechanism of DAG to TAG conversion by DIP2, (ii) the relation between subcellular localisation of DIP2 and its regulatory role in defining the distribution of DAG subspecies at subcellular sites (Baron & Malhotra, 2002; Cowell et al., 2009; Ganesan et al., 2019; Starr & Fratti, 2019; Yang & Kazanietz, 2003), and (iii) how misregulation of a fraction of DAG pool translates into a pathological manifestation.

## Materials and Methods

### Yeast strains and plasmids

All strains used in this study are derivatives of *Saccharomyces cerevisiae* S288C and are listed in **Supplementary Table 2**. Tagging of proteins and gene deletions were performed by standard PCR-based homologous recombination (Longtine et al., 1998). Strains used in this study are isogenic to BY4741 (Mata ura3Δ his3Δ1 leu2Δ0 met15Δ0). Briefly, ScDIP2 (Cmr2; SGD ID: S000005619) gene was replaced with a kanamycin resistance cassette, PCR amplified from pFA6A-KanMX6 plasmid to have flanking sequences homologous to the 5’ and 3’ end of ScDIP2 gene. The PCR product was purified and transformed through direct carrier DNA/PEG method-based transformations (Gietz & Schiestl, 2007) and ScDIP2 deletion mutant colonies (YSM57 and YSM70) were screened for kanamycin selection. Thus, obtained resistant colonies were screened and validated for deletion using PCR through the flanking sequences of kanamycin cassette as primers. YSM101 strain was obtained by C-terminally tagging ScDIP2 at genomic locus with a GFP-KanMX6 cassette amplified from pFA6a-GFP(S65T)-KanMX6 (Longtine et al., 1998). pYSM5 vector was generated by cloning GAL1 promoter and sequence of TEV-mEGFP-8XHis-2XHA in pLE124 vector, a kind gift from Dr. M Palani’s laboratory. ScDIP2 full-length gene and domain-truncated sequences were cloned under GAL1 promoter and tagged C-terminally with TEV-mEGFP-8XHis-2XHA using restriction digestion and ligation cloning method. pYSM10 vector was generated by cloning ScDIP2 sequence into PacI-XhoI sites of pYSM5 vector. pYSM7 vector was generated by cloning ScDIP2 sequence with corresponding promoter and terminator sequence from the genomic locus in pPPM90 vector. Site-directed mutagenesis was performed by PCR using mutagenic oligonucleotides with pYSM10 as the template. Mutagenized plasmids were checked by sequencing. All the primers and plasmids used in this study are in **Supplementary Table 3** and plasmids are listed in **Supplementary Table 4**.

### Media and reagents

Yeast cells were grown at 30°C in a synthetic complete (SC) media. SC media contained 1.7 g/l yeast-nitrogen base with ammonium sulphate (BD Difco™), 20 g/l glucose and was supplemented with the required amino acid mixtures (Sigma) or lacking respective amino acids or supplemented with G418 (200µg/ml) for strains and plasmid selection. For some experiments, yeasts were grown in YP media (Yeast extract-20 g/l, Peptone-20 g/l) supplemented with 2% dextrose or galactose. Different yeast strains were stored at −80°C as glycerol stocks. Prior to all experiments, we briefly thawed the yeast stocks, and few microliters of aliquot was streaked on a SC agar plate and grown under selection at 30°C for 2 days. Before the experiment, single colony was used to prepare seed cultures in 5-10 mL SC media or YP media grown under selection overnight at 30°C in a shaking incubator.

The fly strains used in the study were procured from Bloomington Drosophila Stock Center, Indiana University, USA. DIP2 knock-out fly line from an earlier study (Nitta et.al 2016) was used and was a kind gift from Tetsuya Tabata, University of Tokyo. All stocks and their crosses were maintained in standard cornmeal–yeast–sugar–agar medium at 25°C under a 12–12 h light-dark cycle in the Fly lab, CCMB.

The mouse embryonic stem cells were derived from 3.5 dpc embryo by mating two MmDIP2a^-/-^ mice generated earlier (Kinatukara et al., 2020) in the animal house facility, CCMB using previously described protocols (Behringer et al., 2014). The obtained mouse ES cells were grown as per previously described protocols (Jana et al., 2019). Briefly, cells were cultured on tissue culture-treated plates/dishes coated with 0.1% gelatin, in GMEM media supplemented with L-glutamine, 100 µM β-mercaptoethanol, 1 mM non-essential amino acids (Gibco), 100 units/ml human LIF supplemented with 10% fetal bovine serum (Gibco). Cultures were grown in a humidified incubator at 37°C and 5% CO_2_.

### Metabolic radiolabelling and lipid extraction

Yeast cells were inoculated at 0.05 OD_600_ from an overnight grown primary culture in SC media. 0.5 µCi/ml 1-^14^C acetic acid (American Radiolabeled Chemicals) was added to the media and grown up to mid-log phase (OD_600_ ∼3) to achieve steady-state labelling. Lipids were extracted according to the Folch method (Folch et al., 1957). In brief, equal amounts of cells (10 ml log-phase culture, OD_600_-3.0) were taken and washed with ice-cold 1x phosphate buffered saline (PBS) and then re-suspended in 250 µl of ice-cold PBS. Cell suspension was then transferred to a fresh glass vial and remaining cells in microfuge tube were rinsed with an additional 250 µl of PBS and added to the earlier collection. Acid-washed glass-beads of 0.5 ml volume was added to the cell suspension, followed by the addition of 1.5 ml of chloroform (CHCl_3_)/methanol (MeOH) mix (2:1 ratio). Cells were disrupted by shearing using high speed vortex for 10 min. Phase separation of solvents was achieved by centrifugation at 3000 rpm for 5 min. The organic phase at the bottom part was collected in fresh glass vials. Formic acid (Sigma) was added at 10% v/v to the aqueous phase and vortexed for 1 min. Again, 1 ml of CHCl_3_ was added for re-extraction and vortexed for 5 min, followed by centrifugation and collection of bottom phase. The re-extraction step was repeated 2 times and the pooled organic phase was dried under nitrogen gas at room temperature. Dried samples were stored at −80°C until further use.

### Thin layer chromatography

Dried lipids were resuspended in CHCl_3_: MeOH (1:1) and 10 OD_600_ equivalent lipid was spotted on two separate silica gel TLC plates for neutral and polar lipid separation. Phospholipids were separated using solvent system-A (CHCl_3_: ethanol: water: triethylamine; 30:34:8:35 v/v) (Korte & Casey, 1982). Neutral lipids were separated using solvent system-B (Hexane: diethyl ether: glacial acetic acid; 80:20:2 v/v) (Fakas et al., 2011). Radiolabelled lipids were visualized by a phosphorimager (Amersham Typhoon FLA 9000). Densitometry analysis was performed using ImageJ2 software (Rueden et al., 2017) to measure relative intensities of labelled lipids. The percentages presented for the individual lipid species were normalized to the total ^14^C-labeled fraction.

### Lipidomics sample preparation

The lipid extraction process for different biological samples (Yeast, *Drosophila* and Mouse ES cells) were performed using modified-Folch method as reported previously (Abhyankar et al., 2018; Kelkar et al., 2019; Kumar et al., 2019; Pathak et al., 2018). Briefly, yeast cell pellet harvested from 20 ml culture was washed and suspended in chilled PBS. Cells were disrupted by sonication at 60% amplitude for 1 second ON/ 3 Seconds OFF for 10 cycles with a probe sonicator on ice bed. At this stage, 10 µl of lysates from each sample were collected for protein estimation that is required for data normalization during analysis. Then, CHCl_3_ & MeOH mix with respective internal standards was added to lysate to achieve the ratio of 2:1:1 (CHCl_3_: MeOH: PBS). Mixture was vortexed thoroughly and centrifuged at 3000 rpm at RT to achieve phase separation. Organic bottom phase was collected in fresh glass vials and 100 µl of formic acid (∼ 10% v/v of lysate volume) was added to remaining top phase. After brief vortexing, same volume of CHCl_3_, as mentioned in previous step, was added for re-extraction. Previous steps were repeated to phase separate the mixture, bottom phase was pooled and dried under stream of N_2_ at RT. Protein estimation was performed using Bradford assay reagent (Sigma).

For *Drosophila* samples, 5 days old adult *Drosophila* were collected, flash-frozen in liquid N_2_ and stored at −80°C until further processing. For lipid extraction, 6 flies (3 males and 3 females) were taken and crushed thoroughly in cold PBS with a hand-held homogenizer. Further steps were performed as per the protocol mentioned above.

Mouse ES cells were derived from wild type and MmDIP2A^-/-^ lines as described previously were maintained in GMEM media (supplemented with LIF). ∼ 8 million cells were washed with PBS and flash-frozen in liquid N_2_ before storing at −80°C until further processing. For lipid extraction, Cells were resuspended in ice-cold PBS and disrupted using bath sonicator (1 second ON/ 3 Seconds OFF for 5 cycles). Later the lipidome was extracted as per the protocol mentioned in above.

### Lipidome Analysis by Mass Spectrometry

The extracted lipids were re-solubilized in 200 μL of 2:1 (v/v) CHCl_3_/MeOH, and 20 μL was used for the lipidomics analysis. The extracted lipids were analysed on a Silica F254 thin-layer chromatography and Diethyl Acetate: Hexane: Glacial Acetic Acid (20:80:1) was used as the mobile phase (Hutchins et al., 2008). The lipidomic analysis of the isolated lipid species was carried according to previously described protocols (Abhyankar et al., 2018; Kelkar et al., 2019; Kumar et al., 2019; Pathak et al., 2018). Lipid species were analysed and quantified using the multiple reaction monitoring high resolution (MRM-HR) scanning method on a Sciex X500R QTOF mass spectrometer (MS) fitted with an Exion-LC series UHPLC with minor modifications. All data were acquired and analysed using the SciexOS software. The lipid species were quantified by measuring the area under the curve and the normalized raw MS intensities were used for comparison between various experimental groups. All the lipidomics data are represented as mean ± SEM of six replicates.

### Serial dilution assay

Yeast cells were grown to a final OD_600_ of 0.2 and used as the first dilution. The first dilution stock was used to generate multiple serial dilution stocks (10^-1^, 10^-2^, 10^-3^, and 10^-4^). 5 µl cell suspension from each dilution stocks were spotted sequentially on respective SD agar plates and grown for 48-72 hours at 30°C incubator, before imaging. For different stress assays, chemical reagents were added to nutrient agar media as per the indicated concentrations. For chemical supplementation assay, fatty acids dissolved in DMSO at different molar concentrations were used. Other phospholipid precursors like choline and ethanolamine were dissolved in autoclaved water and used for supplementation assays.

### UPR reporter assay

pPM47 (UPR-RFP CEN/ARS URA3) was a gift from Feroz Papa (Addgene plasmid # 20132). The experiment was performed according to the procedures described earlier (Merksamer et al., 2008). Briefly, cells harbouring single-copy reporter plasmid was grown in respective media up to early-log, mid-log phase or stationary phase, harvested by centrifugation and washed with PBS before experiment. Fluorescence signal was measured with a flow cytometer, BD LSR Fortessa (excitation-561 nm, emission-610 nm, volume-100 µL, number of cells-10000). Cells without reporter plasmid served as a control. To validate the UPR reporter assay, UPR signal was measured using cultures with increasing concentration of tunicamycin added and incubated for 2 hours at 30°C with shaking. For chemical supplementation, oleic acid was added to media at 1mM concentration and grown up to log phase before subjecting for fluorescence measurement. Data analysis was performed using FCS Express (De novo software).

### Growth curve and cell viability assay

The growth curve was obtained by growing cells from an equal initial OD_600_ of 0.2. Cultures were allowed to grow in 30°C incubator shaker and the OD_600_ was recorded every 4 hours. These experiments were repeated in triplicates and the error bars represent standard deviation values.

For hypertonic stress cell viability assay, overnight grown cells were diluted to OD_600_ of 0.2 in SC-dextrose media and grown for 4-5 hours in 30°C incubator shaker. Log phase cells were harvested, diluted to OD_600_ of 0.2 and grown in presence of 1 M NaCl in YP-dextrose media. After 4-5 hours of growth, cells were incubated with 0.2% (v/v) trypan blue (Sigma) for 5 min and further washed five times with PBS. Cell viability was assessed using bright field microscopy by counting the proportion of live cells (unstained) to dead cells (stained). Cells grown without NaCl was used as control.

For hypotonic stress cell viability assay, overnight grown primary cultures were diluted in SC-galactose media at OD_600_ of 0.2. After growing till log phase, cells were diluted in 5 ml YP-galactose media with 1 M NaCl and incubated for 15 minutes at 30°C with 200 rpm shaking. Cells were harvested by centrifugation and re-dispersed in 5 ml autoclaved filtered Milli-Q (hypotonic condition), After incubation for 15 minutes at 30°C with 200 rpm shaking, cell death was assessed as per the protocol mentioned above. Cells incubated without salt and hypotonic condition were used as control.

### Live cell microscopy

ScDIP2-GFP knock-in strain was grown overnight in 5 ml of SC broth at 30°C. 20 nM of MitoTracker™ Red CMXRos (Invitrogen Inc.) was added to these overnight grown cells and incubated further for 30 minutes with shaking. The MitoTracker™ incubated cells were then washed in autoclaved water. 5 µl of those cells were immobilized on a Poly-*L*-lysine coated glass slides and visualized under a Zeiss Axio Imager.Z2, inverted widefield fluorescence microscope equipped with a HAL 100 illuminator, Plan Apochromat 100X oil objective (NA 1.4), and a AxioCam CCD camera. Images were captured in DIC (Nomarski optics), Texas red (for MitoTracker™ Red) and eGFP fluorescence mode. The vacuole of ScDIP2-GFP knock-in cells were stained with FM4-64 dye (Invitrogen) following the protocol described in ‘Vacuole morphology assay’ section. Cells were further processed, and images were collected in same procedure described above. All the experiments were done in biological triplicates.

Sec63-mRFP (pSM1960, a gift from Susan Michaelis; Addgene plasmid # 41842) (Metzger et al., 2008) was used as ER marker. For nuclear staining, cells were fixed with 1% formaldehyde followed by incubation with 1 μg/ml DAPI (Sigma) in PBS for 10 min without shaking and subsequent washing with PBS. All images were processed using ImageJ2 software.

### Vacuole morphology assay

Vacuole morphology was determined by FM4-64 staining of the vacuole membrane as described previously (Vida & Emr, 1995). Briefly, overnight grown culture was inoculated in a fresh medium with and grown up to early-log phase (OD_600_-1.2 -1.5). Cells equivalent to 1.5 OD_600_ were harvested by spinning at 4000 rpm for 5 minutes, washed with PBS and resuspended in 200 µl of media. Cells were pulse-labelled with 10 µM of FM4-64 for 30 minutes at 30°C, then washed in medium without dye and finally suspended in 5ml media to grow for 90 minutes at 30°C, 200 rpm. Cells were then harvested by centrifugation, washed twice with PBS and resuspended in PBS or SC media for microscopy. Diameter of the largest vacuole of budding yeast cell was measured manually using the line tool of ImageJ. Vacuolar diameters for indicated strains were plotted into box and whisker plots.

For vacuole fission-fusion assay (Jones et al., 2010), yeast vacuoles were stained first according to the above-mentioned protocol. It is now well established that hypotonic stress induces vacuole fusion while hypertonic stress induces vacuole fission in vivo. Therefore, cells were subjected to the hypertonic condition, i.e., YP-dextrose or YP-galactose with 0.4 M NaCl and grown for 10-15 minutes at 30°C, 200 rpm. The vacuole fission was confirmed by visualizing under fluorescence microscope. To induce vacuole fusion, cells were harvested and subjected to hypotonic condition, i.e., 10-fold dilution in autoclaved Milli-Q water and grown for 10 minutes, 30°C, shaking at 200 rpm. Images were captured by putting washed cells on 2% low-melting agar bed on glass slides and mounted with a cover slip.

### Western blot analysis

For GFP-tagged full-length and domain-truncated constructs of ScDIP2 expressing strains, the cell pellet was re-suspended in 20% trichloroacetic acid (TCA), mixed with ice cold glass beads (0.5 mm diameter) and ruptured by bead beating for 3 cycles (1-minute ON,1-minute OFF). The supernatant was collected, and the pellet was harvested by centrifugation at 14000 rpm for 10 minutes. Pellet was washed with ice-cold acetone, then dried under vacuum and finally dissolved in SDS-loading buffer. All the steps were performed at 4°C. Protein extracts were separated by Sodium dodecyl sulphate-polyacrylamide gel electrophoresis (SDS-PAGE). After separation, proteins were transferred onto a nitrocellulose membrane by wet transfer method at 100V for 2 hours. Membranes were blocked with 5% non-fat milk powder in TBST buffer (20 mM Tris-HCl pH-7.5, 150 mM NaCl, 0.1% Tween-20) and immunoblotting was performed using anti-Pgk1 (monoclonal, Thermo Fisher Scientific) and anti-GFP (monoclonal, Cell Signaling Technologies). Signals were developed using the BioRad ChemiDoc MP imaging system.

### Isolation of total membrane

Total endomembrane fraction was isolated as described previously (Sorin et al., 1997). ScDIP2-GFP knock-in cells were resuspended in a cell resuspension buffer (50 mM Tris/HCl, pH 7.6, 0.6 M sorbitol, 1mM EDTA) containing protease inhibitor cocktail (Calbiochem) and 1mM PMSF (Sigma). Cells were disrupted by vortexing the cell suspension with acid-washed chilled glass beads for 8 cycles of 30 seconds pulse and 30 seconds chilling on ice. Lysate was centrifuged at 1000g for 10 minutes to separate cell debris as a pellet and collect the supernatant that contains organelles, membranes and cytoplasmic proteins. The crude membrane fraction was collected by centrifuging the supernatant at 100000g for 1-hour at 4°C and further resuspending the pellet in membrane resuspension buffer (20 mM Tris/HCL, 0.3 M sucrose and 0.1 mM CaCl_2_). The fractions were subjected to SDS-PAGE and ScDIP2-GFP was probed with anti-GFP antibody as per protocol mentioned in above section.

### Bioinformatics analysis

All sequences were identified and retrieved from the NCBI sequence database using ScDIP2 (UNIPROT ID: Q12275) as a template and BLAST search algorithm (Altschul et al., 1990). Domain boundaries of ScDIP2 were marked based on consensus obtained from structure-based sequence alignments generated in EXPRESSO (Armougom et al., 2006), manual inspection of homology models, generated via MODELLER (Webb & Sali, 2016), visualized in PyMOL (Schrödinger, 2015) and domain definitions obtained from Conserved Domain Database (Lu et al., 2020). Multiple sequence alignment was generated using MAFFT (Katoh et al., 2002) and MUSCLE (Edgar, 2004). The sequence alignments were rendered using ESPript 3.0 (Robert & Gouet, 2014). The phylogenetic analysis of these sequences was carried out using maximum-likelihood method as implemented in IQTREE (Nguyen et al., 2015) using default parameters. The phylogenetic tree was rendered in iTOL server (Letunic & Bork, 2021). The images of various organisms were reusable silhouette images of organisms obtained from PhyloPic (www.phylopic.org), under Creative Commons license. Pairwise sequence identity was calculated using SIAS (http://imed.med.ucm.es/Tools/sias.html) with default parameters. Sequence logos were generated using WebLogo (Crooks et al., 2004). Homology models for structural analysis were obtained from AlphaFold Protein Structure Database (Jumper et al., 2021; Varadi et al., 2021).

The GO terms were tabulated by analysing the genetic interactions using DRYGIN database (https://thecellmap.org/) (Koh et al., 2010) at default stringency levels (between + 0.08 and - 0.08) along with tools from YeastMine (Balakrishnan et al., 2012) and Gene Ontology Slim Term Mapper (Ashburner et al., 2000).

### Statistics

Error bars represent standard error of the mean (SEM) or standard deviation (SD) as indicated in the respective figure legends. Data were processed and analysed using Microsoft Excel and GraphPad Prism v.8.0.2.263. Statistical analysis of the differences between two groups was performed using a two-tailed, unpaired t-test. Based on the p-value obtained, the significance of differences marked as *, p < 0.05; **, p < 0.01; ***, p < 0.001; ****, p<0.0001; ns, not significant.

## Supporting information

Supplementary Figures with legends

## Acknowledgments

We thank Palani Murugan Rangasamy (CSIR-CCMB) for helping in generation of knock-out strains of yeast and for sharing plasmids; Animal house facility, Fly lab facility, Advanced Microscopy facility and Flow cytometry (FACS) facility for help in performing experiments. We thank Krishnaveni Mishra (UoH) for sharing yeast knock-out strains and Rakesh Mishra (CSIR-CCMB) for helping with *Drosophila* experiments. We thank Koustav Sanyal (JNCASR) and Hashim Reza (JNCASR) for insightful discussion on vacuole morphology; Rajesh Gokhale (NII), Durgadas P Kasbekar (CDFD) and Venkat R Chalamcharla (CSIR-CCMB) for their valuable comments on the manuscript. SM thank CSIR, India and PK thank the Department of Biotechnology, India for research fellowship. SaS thanks UGC, India for research fellowship. SSK thank DBT/Wellcome Trust India Alliance Fellowship (grant number IA/I/15/2/502058) and a Department of Science and Technology (DST) Fund for Improvement of S&T Infrastructure (grant number SR/FST/LSII-043/2016) to the IISER Pune Biology Department. RS thank NCP under health care theme project of CSIR, India; J.C. Bose Fellowship of SERB, India; and Centre of Excellence Project of Department of Biotechnology, India.

## Authors contribution

SM and PK designed and performed the experiments, analysed and interpreted data. SaS, GP, ND and BP helped with cloning plasmids, microscopy, and western blot experiments. SSK designed the lipidomics experiments. ShuS and SSK performed LC/MS experiments and data analysis. SHS generated and maintained the *Drosophila* lines and CSP derived ES cells from mouse knock-out lines. RSN conceived and supervised the study. SM, PK and RSN wrote the manuscript with the help of SSK and BP. All the authors have analysed and reviewed the data and manuscript.

## References

Abhyankar, V., Kaduskar, B., Kamat, S. S., Deobagkar, D., & Ratnaparkhi, G. S. (2018). Drosophila DNA/RNA methyltransferase contributes to robust host defense in aging animals by regulating sphingolipid metabolism. J Exp Biol, 221(Pt 22). https://doi.org/10.1242/jeb.187989

Altschul, S. F., Gish, W., Miller, W., Myers, E. W., & Lipman, D. J. (1990). Basic local alignment search tool. J Mol Biol, 215(3), 403–410. https://doi.org/10.1016/S0022-2836(05)80360-2

Armougom, F., Moretti, S., Poirot, O., Audic, S., Dumas, P., Schaeli, B., Keduas, V., & Notredame, C. (2006). Expresso: automatic incorporation of structural information in multiple sequence alignments using 3D-Coffee. Nucleic Acids Res, 34(Web Server issue), W604–608. https://doi.org/10.1093/nar/gkl092

Arora, P., Goyal, A., Natarajan, V. T., Rajakumara, E., Verma, P., Gupta, R., Yousuf, M., Trivedi, O. A., Mohanty, D., Tyagi, A., Sankaranarayanan, R., & Gokhale, R. S. (2009). Mechanistic and functional insights into fatty acid activation in Mycobacterium tuberculosis. Nat Chem Biol, 5(3), 166–173. https://doi.org/10.1038/nchembio.143

Ashburner, M., Ball, C. A., Blake, J. A., Botstein, D., Butler, H., Cherry, J. M., Davis, A. P., Dolinski, K., Dwight, S. S., Eppig, J. T., Harris, M. A., Hill, D. P., Issel-Tarver, L., Kasarskis, A., Lewis, S., Matese, J. C., Richardson, J. E., Ringwald, M., Rubin, G. M., & Sherlock, G. (2000). Gene ontology: tool for the unification of biology. The Gene Ontology Consortium. Nat Genet, 25(1), 25–29. https://doi.org/10.1038/75556

Atilla-Gokcumen, G. E., Muro, E., Relat-Goberna, J., Sasse, S., Bedigian, A., Coughlin, M. L., Garcia-Manyes, S., & Eggert, U. S. (2014). Dividing cells regulate their lipid composition and localization. Cell, 156(3), 428–439. https://doi.org/10.1016/j.cell.2013.12.015

Balakrishnan, R., Park, J., Karra, K., Hitz, B. C., Binkley, G., Hong, E. L., Sullivan, J., Micklem, G., & Cherry, J. M. (2012). YeastMine--an integrated data warehouse for Saccharomyces cerevisiae data as a multipurpose tool-kit. Database (Oxford), 2012, bar062. https://doi.org/10.1093/database/bar062

Baron, C. L., & Malhotra, V. (2002). Role of diacylglycerol in PKD recruitment to the TGN and protein transport to the plasma membrane. Science, 295(5553), 325–328. https://doi.org/10.1126/science.1066759

Behringer, R., Gertsenstein, M., Nagy, K. V., & Nagy, A. (2014). Manipulating the Mouse Embryo: A Laboratory Manual. Cold Spring Harbor Laboratory Press. https://books.google.ne/books?id=LR2anQEACAAJ

Bigay, J., & Antonny, B. (2012). Curvature, lipid packing, and electrostatics of membrane organelles: defining cellular territories in determining specificity. Dev Cell, 23(5), 886–895. https://doi.org/10.1016/j.devcel.2012.10.009

Boumann, H. A., de Kruijff, B., Heck, A. J., & de Kroon, A. I. (2004). The selective utilization of substrates in vivo by the phosphatidylethanolamine and phosphatidylcholine biosynthetic enzymes Ept1p and Cpt1p in yeast. FEBS Lett, 569(1-3), 173–177. https://doi.org/10.1016/j.febslet.2004.05.043

Brachmann, C. B., Davies, A., Cost, G. J., Caputo, E., Li, J., Hieter, P., & Boeke, J. D. (1998). Designer deletion strains derived from Saccharomyces cerevisiae S288C: a useful set of strains and plasmids for PCR-mediated gene disruption and other applications. Yeast, 14(2), 115–132. https://doi.org/10.1002/(SICI)1097-0061(19980130)14:2<115::AID-YEA204>3.0.CO;2-2

Carrasco, S., & Merida, I. (2007). Diacylglycerol, when simplicity becomes complex. Trends Biochem Sci, 32(1), 27–36. https://doi.org/10.1016/j.tibs.2006.11.004

Casanovas, A., Sprenger, R. R., Tarasov, K., Ruckerbauer, D. E., Hannibal-Bach, H. K., Zanghellini, J., Jensen, O. N., & Ejsing, C. S. (2015). Quantitative analysis of proteome and lipidome dynamics reveals functional regulation of global lipid metabolism. Chem Biol, 22(3), 412–425. https://doi.org/10.1016/j.chembiol.2015.02.007

Chen, Y., Feldman, D. E., Deng, C., Brown, J. A., De Giacomo, A. F., Gaw, A. F., Shi, G., Le, Q. T., Brown, J. M., & Koong, A. C. (2005). Identification of mitogen-activated protein kinase signaling pathways that confer resistance to endoplasmic reticulum stress in Saccharomyces cerevisiae. Mol Cancer Res, 3(12), 669–677. https://doi.org/10.1158/1541-7786.MCR-05-0181

Cowell, C. F., Doppler, H., Yan, I. K., Hausser, A., Umezawa, Y., & Storz, P. (2009). Mitochondrial diacylglycerol initiates protein-kinase D1-mediated ROS signaling. J Cell Sci, 122(Pt 7), 919–928. https://doi.org/10.1242/jcs.041061

Crooks, G. E., Hon, G., Chandonia, J. M., & Brenner, S. E. (2004). WebLogo: a sequence logo generator. Genome Res, 14(6), 1188–1190. https://doi.org/10.1101/gr.849004

Edgar, R. C. (2004). MUSCLE: multiple sequence alignment with high accuracy and high throughput. Nucleic Acids Res, 32(5), 1792–1797. https://doi.org/10.1093/nar/gkh340

Egger, G., Roetzer, K. M., Noor, A., Lionel, A. C., Mahmood, H., Schwarzbraun, T., Boright, O., Mikhailov, A., Marshall, C. R., Windpassinger, C., Petek, E., Scherer, S. W., Kaschnitz, W., & Vincent, J. B. (2014). Identification of risk genes for autism spectrum disorder through copy number variation analysis in Austrian families. Neurogenetics, 15(2), 117–127. https://doi.org/10.1007/s10048-014-0394-0

Ejsing, C. S., Sampaio, J. L., Surendranath, V., Duchoslav, E., Ekroos, K., Klemm, R. W., Simons, K., & Shevchenko, A. (2009). Global analysis of the yeast lipidome by quantitative shotgun mass spectrometry. Proc Natl Acad Sci U S A, 106(7), 2136–2141. https://doi.org/10.1073/pnas.0811700106

Fakas, S., Konstantinou, C., & Carman, G. M. (2011). DGK1-encoded diacylglycerol kinase activity is required for phospholipid synthesis during growth resumption from stationary phase in Saccharomyces cerevisiae. J Biol Chem, 286(2), 1464–1474. https://doi.org/10.1074/jbc.M110.194308

Folch, J., Lees, M., & Sloane Stanley, G. H. (1957). A simple method for the isolation and purification of total lipides from animal tissues. J Biol Chem, 226(1), 497–509. https://www.ncbi.nlm.nih.gov/pubmed/13428781

Ganesan, S., Sosa Ponce, M. L., Tavassoli, M., Shabits, B. N., Mahadeo, M., Prenner, E. J., Terebiznik, M. R., & Zaremberg, V. (2019). Metabolic control of cytosolic-facing pools of diacylglycerol in budding yeast. Traffic, 20(3), 226–245. https://doi.org/10.1111/tra.12632

Gietz, R. D., & Schiestl, R. H. (2007). High-efficiency yeast transformation using the LiAc/SS carrier DNA/PEG method. Nat Protoc, 2(1), 31–34. https://doi.org/10.1038/nprot.2007.13

Gilbert, J., & Man, H. Y. (2017). Fundamental Elements in Autism: From Neurogenesis and Neurite Growth to Synaptic Plasticity. Front Cell Neurosci, 11, 359. https://doi.org/10.3389/fncel.2017.00359

Gong, J., Qiu, C., Huang, D., Zhang, Y., Yu, S., & Zeng, C. (2018). Integrative functional analysis of super enhancer SNPs for coronary artery disease. J Hum Genet, 63(5), 627–638. https://doi.org/10.1038/s10038-018-0422-2

Goyal, A., Verma, P., Anandhakrishnan, M., Gokhale, R. S., & Sankaranarayanan, R. (2012). Molecular basis of the functional divergence of fatty acyl-AMP ligase biosynthetic enzymes of Mycobacterium tuberculosis. J Mol Biol, 416(2), 221–238. https://doi.org/10.1016/j.jmb.2011.12.031

Grevengoed, T. J., Klett, E. L., & Coleman, R. A. (2014). Acyl-CoA metabolism and partitioning. Annu Rev Nutr, 34, 1–30. https://doi.org/10.1146/annurev-nutr-071813-105541

Gulick, A. M. (2009). Conformational dynamics in the Acyl-CoA synthetases, adenylation domains of non-ribosomal peptide synthetases, and firefly luciferase. ACS Chem Biol, 4(10), 811–827. https://doi.org/10.1021/cb900156h

Han, S., Lone, M. A., Schneiter, R., & Chang, A. (2010). Orm1 and Orm2 are conserved endoplasmic reticulum membrane proteins regulating lipid homeostasis and protein quality control. Proc Natl Acad Sci U S A, 107(13), 5851–5856. https://doi.org/10.1073/pnas.0911617107

Harayama, T., & Riezman, H. (2018). Understanding the diversity of membrane lipid composition. Nat Rev Mol Cell Biol, 19(5), 281–296. https://doi.org/10.1038/nrm.2017.138

Heier, C., & Kuhnlein, R. P. (2018). Triacylglycerol Metabolism in Drosophila melanogaster. Genetics, 210(4), 1163–1184. https://doi.org/10.1534/genetics.118.301583

Henneberry, A. L., Lagace, T. A., Ridgway, N. D., & McMaster, C. R. (2001). Phosphatidylcholine synthesis influences the diacylglycerol homeostasis required for SEC14p-dependent Golgi function and cell growth. Mol Biol Cell, 12(3), 511–520. https://doi.org/10.1091/mbc.12.3.511

Hutchins, P. M., Barkley, R. M., & Murphy, R. C. (2008). Separation of cellular nonpolar neutral lipids by normal-phase chromatography and analysis by electrospray ionization mass spectrometry. J Lipid Res, 49(4), 804–813. https://doi.org/10.1194/jlr.M700521-JLR200

Iossifov, I., Ronemus, M., Levy, D., Wang, Z., Hakker, I., Rosenbaum, J., Yamrom, B., Lee, Y. H., Narzisi, G., Leotta, A., Kendall, J., Grabowska, E., Ma, B., Marks, S., Rodgers, L., Stepansky, A., Troge, J., Andrews, P., Bekritsky, M., … Wigler, M. (2012). De novo gene disruptions in children on the autistic spectrum. Neuron, 74(2), 285–299. https://doi.org/10.1016/j.neuron.2012.04.009

Jacquier, N., Choudhary, V., Mari, M., Toulmay, A., Reggiori, F., & Schneiter, R. (2011). Lipid droplets are functionally connected to the endoplasmic reticulum in Saccharomyces cerevisiae. J Cell Sci, 124(Pt 14), 2424–2437. https://doi.org/10.1242/jcs.076836

Jana, D., Kale, H. T., & Shekar, P. C. (2019). Generation of Cdx2-mCherry knock-in murine ES cell line to model trophectoderm and intestinal lineage differentiation. Stem Cell Res, 39, 101521. https://doi.org/10.1016/j.scr.2019.101521

Jiao, X., Wood, L. D., Lindman, M., Jones, S., Buckhaults, P., Polyak, K., Sukumar, S., Carter, H., Kim, D., Karchin, R., & Sjoblom, T. (2012). Somatic mutations in the Notch, NF-KB, PIK3CA, and Hedgehog pathways in human breast cancers. Genes Chromosomes Cancer, 51(5), 480–489. https://doi.org/10.1002/gcc.21935

Jones, L., Tedrick, K., Baier, A., Logan, M. R., & Eitzen, G. (2010). Cdc42p is activated during vacuole membrane fusion in a sterol-dependent subreaction of priming. J Biol Chem, 285(7), 4298–4306. https://doi.org/10.1074/jbc.M109.074609

Jonikas, M. C., Collins, S. R., Denic, V., Oh, E., Quan, E. M., Schmid, V., Weibezahn, J., Schwappach, B., Walter, P., Weissman, J. S., & Schuldiner, M. (2009). Comprehensive characterization of genes required for protein folding in the endoplasmic reticulum. Science, 323(5922), 1693–1697. https://doi.org/10.1126/science.1167983

Joshi, A. S., Nebenfuehr, B., Choudhary, V., Satpute-Krishnan, P., Levine, T. P., Golden, A., & Prinz, W. A. (2018). Lipid droplet and peroxisome biogenesis occur at the same ER subdomains. Nat Commun, 9(1), 2940. https://doi.org/10.1038/s41467-018-05277-3

Jumper, J., Evans, R., Pritzel, A., Green, T., Figurnov, M., Ronneberger, O., Tunyasuvunakool, K., Bates, R., Zidek, A., Potapenko, A., Bridgland, A., Meyer, C., Kohl, S. A. A., Ballard, A. J., Cowie, A., Romera-Paredes, B., Nikolov, S., Jain, R., Adler, J., … Hassabis, D. (2021). Highly accurate protein structure prediction with AlphaFold. Nature, 596(7873), 583–589. https://doi.org/10.1038/s41586-021-03819-2

Katoh, K., Misawa, K., Kuma, K., & Miyata, T. (2002). MAFFT: a novel method for rapid multiple sequence alignment based on fast Fourier transform. Nucleic Acids Res, 30(14), 3059–3066. https://doi.org/10.1093/nar/gkf436

Kelkar, D. S., Ravikumar, G., Mehendale, N., Singh, S., Joshi, A., Sharma, A. K., Mhetre, A., Rajendran, A., Chakrapani, H., & Kamat, S. S. (2019). A chemical-genetic screen identifies ABHD12 as an oxidized-phosphatidylserine lipase. Nat Chem Biol, 15(2), 169–178. https://doi.org/10.1038/s41589-018-0195-0

Khandelwal, N., Shaikh, M., Mhetre, A., Singh, S., Sajeevan, T., Joshi, A., Balaji, K. N., Chakrapani, H., & Kamat, S. S. (2021). Fatty acid chain length drives lysophosphatidylserine-dependent immunological outputs. Cell Chem Biol. https://doi.org/10.1016/j.chembiol.2021.01.008

Kinatukara, P., Subramaniyan, P. S., Patil, G. S., Shambhavi, S., Singh, S., Mhetre, A., Madduri, M. K., Soundararajan, A., Patel, K. D., Shekar, P. C., Kamat, S. S., Kumar, S., & Sankaranarayanan, R. (2020). Peri-natal growth retardation rate and fat mass accumulation in mice lacking Dip2A is dependent on the dietary composition. Transgenic Res, 29(5-6), 553–562. https://doi.org/10.1007/s11248-020-00219-6

Koh, J. L., Ding, H., Costanzo, M., Baryshnikova, A., Toufighi, K., Bader, G. D., Myers, C. L., Andrews, B. J., & Boone, C. (2010). DRYGIN: a database of quantitative genetic interaction networks in yeast. Nucleic Acids Res, 38(Database issue), D502-507. https://doi.org/10.1093/nar/gkp820

Kong, R., Shao, S., Wang, J., Zhang, X., Guo, S., Zou, L., Zhong, R., Lou, J., Zhou, J., Zhang, J., & Song, R. (2016). Genetic variant in DIP2A gene is associated with developmental dyslexia in Chinese population. Am J Med Genet B Neuropsychiatr Genet, 171B(2), 203-208. https://doi.org/10.1002/ajmg.b.32392

Korte, K., & Casey, M. L. (1982). Phospholipid and neutral lipid separation by one-dimensional thin-layer chromatography. J Chromatogr, 232(1), 47–53. https://doi.org/10.1016/s0378-4347(00)86006-5

Kumar, M., Ojha, S., Rai, P., Joshi, A., Kamat, S. S., & Mallik, R. (2019). Insulin activates intracellular transport of lipid droplets to release triglycerides from the liver. J Cell Biol, 218(11), 3697–3713. https://doi.org/10.1083/jcb.201903102

Larsson, C., Ali, M. A., Pandzic, T., Lindroth, A. M., He, L., & Sjoblom, T. (2017). Loss of DIP2C in RKO cells stimulates changes in DNA methylation and epithelial-mesenchymal transition. BMC Cancer, 17(1), 487. https://doi.org/10.1186/s12885-017-3472-5

Lee, C., Fisher, S. K., Agranoff, B. W., & Hajra, A. K. (1991). Quantitative analysis of molecular species of diacylglycerol and phosphatidate formed upon muscarinic receptor activation of human SK-N-SH neuroblastoma cells. J Biol Chem, 266(34), 22837–22846. https://www.ncbi.nlm.nih.gov/pubmed/1744076

Letunic, I., & Bork, P. (2021). Interactive Tree Of Life (iTOL) v5: an online tool for phylogenetic tree display and annotation. Nucleic Acids Res, 49(W1), W293–W296. https://doi.org/10.1093/nar/gkab301

Li, D., Yang, S. G., He, C. W., Zhang, Z. T., Liang, Y., Li, H., Zhu, J., Su, X., Gong, Q., & Xie, Z. (2020). Excess diacylglycerol at the endoplasmic reticulum disrupts endomembrane homeostasis and autophagy. BMC Biol, 18(1), 107. https://doi.org/10.1186/s12915-020-00837-w

Longtine, M. S., McKenzie, A., 3rd, Demarini, D. J., Shah, N. G., Wach, A., Brachat, A., Philippsen, P., & Pringle, J. R. (1998). Additional modules for versatile and economical PCR-based gene deletion and modification in Saccharomyces cerevisiae. Yeast, 14(10), 953-961. https://doi.org/10.1002/(SICI)1097-0061(199807)14:10<953::AID-YEA293>3.0.CO;2-U

Lu, S., Wang, J., Chitsaz, F., Derbyshire, M. K., Geer, R. C., Gonzales, N. R., Gwadz, M., Hurwitz, D. I., Marchler, G. H., Song, J. S., Thanki, N., Yamashita, R. A., Yang, M., Zhang, D., Zheng, C., Lanczycki, C. J., & Marchler-Bauer, A. (2020). CDD/SPARCLE: the conserved domain database in 2020. Nucleic Acids Res, 48(D1), D265–D268. https://doi.org/10.1093/nar/gkz991

Lu, S. W., Kroken, S., Lee, B. N., Robbertse, B., Churchill, A. C., Yoder, O. C., & Turgeon, B. G. (2003). A novel class of gene controlling virulence in plant pathogenic ascomycete fungi. Proc Natl Acad Sci U S A, 100(10), 5980–5985. https://doi.org/10.1073/pnas.0931375100

Ma, J., Zhang, L. Q., He, Z. X., He, X. X., Wang, Y. J., Jian, Y. L., Wang, X., Zhang, B. B., Su, C., Lu, J., Huang, B. Q., Zhang, Y., Wang, G. Y., Guo, W. X., Qiu, D. L., Mei, L., Xiong, W. C., Zheng, Y. W., & Zhu, X. J. (2019). Autism candidate gene DIP2A regulates spine morphogenesis via acetylation of cortactin. PLoS Biol, 17(10), e3000461. https://doi.org/10.1371/journal.pbio.3000461

Marignani, P. A., Epand, R. M., & Sebaldt, R. J. (1996). Acyl chain dependence of diacylglycerol activation of protein kinase C activity in vitro. Biochem Biophys Res Commun, 225(2), 469–473. https://doi.org/10.1006/bbrc.1996.1196

Markgraf, D. F., Klemm, R. W., Junker, M., Hannibal-Bach, H. K., Ejsing, C. S., & Rapoport, T. A. (2014). An ER protein functionally couples neutral lipid metabolism on lipid droplets to membrane lipid synthesis in the ER. Cell Rep, 6(1), 44–55. https://doi.org/10.1016/j.celrep.2013.11.046

Merksamer, P. I., Trusina, A., & Papa, F. R. (2008). Real-time redox measurements during endoplasmic reticulum stress reveal interlinked protein folding functions. Cell, 135(5), 933–947. https://doi.org/10.1016/j.cell.2008.10.011

Metzger, M. B., Maurer, M. J., Dancy, B. M., & Michaelis, S. (2008). Degradation of a cytosolic protein requires endoplasmic reticulum-associated degradation machinery. J Biol Chem, 283(47), 32302–32316. https://doi.org/10.1074/jbc.M806424200

Milne, S. B., Ivanova, P. T., Armstrong, M. D., Myers, D. S., Lubarda, J., Shulga, Y. V., Topham, M. K., Brown, H. A., & Epand, R. M. (2008). Dramatic differences in the roles in lipid metabolism of two isoforms of diacylglycerol kinase. Biochemistry, 47(36), 9372–9379. https://doi.org/10.1021/bi800492c

Montell, E., Turini, M., Marotta, M., Roberts, M., Noe, V., Ciudad, C. J., Mace, K., & Gomez-Foix, A. M. (2001). DAG accumulation from saturated fatty acids desensitizes insulin stimulation of glucose uptake in muscle cells. Am J Physiol Endocrinol Metab, 280(2), E229–237. https://doi.org/10.1152/ajpendo.2001.280.2.E229

Mora, G., Scharnewski, M., & Fulda, M. (2012). Neutral lipid metabolism influences phospholipid synthesis and deacylation in Saccharomyces cerevisiae. PLoS One, 7(11), e49269. https://doi.org/10.1371/journal.pone.0049269

Narra, H. P., Shubitz, L. F., Mandel, M. A., Trinh, H. T., Griffin, K., Buntzman, A. S., Frelinger, J. A., Galgiani, J. N., & Orbach, M. J. (2016). A Coccidioides posadasii CPS1 Deletion Mutant Is Avirulent and Protects Mice from Lethal Infection. Infect Immun, 84(10), 3007–3016. https://doi.org/10.1128/IAI.00633-16

Nguyen, L. T., Schmidt, H. A., von Haeseler, A., & Minh, B. Q. (2015). IQ-TREE: a fast and effective stochastic algorithm for estimating maximum-likelihood phylogenies. Mol Biol Evol, 32(1), 268–274. https://doi.org/10.1093/molbev/msu300

Nitta, Y., Yamazaki, D., Sugie, A., Hiroi, M., & Tabata, T. (2017). DISCO Interacting Protein 2 regulates axonal bifurcation and guidance of Drosophila mushroom body neurons. Dev Biol, 421(2), 233–244. https://doi.org/10.1016/j.ydbio.2016.11.015

Noblett, N., Wu, Z., Ding, Z. H., Park, S., Roenspies, T., Flibotte, S., Chisholm, A. D., Jin, Y., & Colavita, A. (2019). DIP-2 suppresses ectopic neurite sprouting and axonal regeneration in mature neurons. J Cell Biol, 218(1), 125–133. https://doi.org/10.1083/jcb.201804207

Oelkers, P., Cromley, D., Padamsee, M., Billheimer, J. T., & Sturley, S. L. (2002). The DGA1 gene determines a second triglyceride synthetic pathway in yeast. J Biol Chem, 277(11), 8877–8881. https://doi.org/10.1074/jbc.M111646200

Oelkers, P., Tinkelenberg, A., Erdeniz, N., Cromley, D., Billheimer, J. T., & Sturley, S. L. (2000). A lecithin cholesterol acyltransferase-like gene mediates diacylglycerol esterification in yeast. J Biol Chem, 275(21), 15609–15612. https://doi.org/10.1074/jbc.C000144200

Pathak, D., Mehendale, N., Singh, S., Mallik, R., & Kamat, S. S. (2018). Lipidomics Suggests a New Role for Ceramide Synthase in Phagocytosis. ACS Chem Biol, 13(8), 2280–2287. https://doi.org/10.1021/acschembio.8b00438

Patil, G. S., Kinatukara, P., Mondal, S., Shambhavi, S., Patel, K. D., Pramanik, S., Dubey, N., Narasimhan, S., Madduri, M. K., Pal, B., Gokhale, R. S., & Sankaranarayanan, R. (2021). A universal pocket in Fatty acyl-AMP ligases ensures redirection of fatty acid pool away from Coenzyme A-based activation. Elife, 10. https://doi.org/10.7554/eLife.70067

Phillips, M., & Pozzo-Miller, L. (2015). Dendritic spine dysgenesis in autism related disorders. Neurosci Lett, 601, 30–40. https://doi.org/10.1016/j.neulet.2015.01.011

Poelmans, G., Engelen, J. J., Van Lent-Albrechts, J., Smeets, H. J., Schoenmakers, E., Franke, B., Buitelaar, J. K., Wuisman-Frerker, M., Erens, W., Steyaert, J., & Schrander-Stumpel, C. (2009). Identification of novel dyslexia candidate genes through the analysis of a chromosomal deletion. Am J Med Genet B Neuropsychiatr Genet, 150B(1), 140-147. https://doi.org/10.1002/ajmg.b.30787

Raghu, P. (2020). Functional diversity in a lipidome. Proc Natl Acad Sci U S A, 117(21), 11191–11193. https://doi.org/10.1073/pnas.2004764117

Robert, X., & Gouet, P. (2014). Deciphering key features in protein structures with the new ENDscript server. Nucleic Acids Res, 42(Web Server issue), W320–324. https://doi.org/10.1093/nar/gku316

Rockenfeller, P., Smolnig, M., Diessl, J., Bashir, M., Schmiedhofer, V., Knittelfelder, O., Ring, J., Franz, J., Foessl, I., Khan, M. J., Rost, R., Graier, W. F., Kroemer, G., Zimmermann, A., Carmona-Gutierrez, D., Eisenberg, T., Buttner, S., Sigrist, S. J., Kuhnlein, R. P., … Madeo, F. (2018). Diacylglycerol triggers Rim101 pathway-dependent necrosis in yeast: a model for lipotoxicity. Cell Death Differ, 25(4), 767–783. https://doi.org/10.1038/s41418-017-0014-2

Ron, D., & Walter, P. (2007). Signal integration in the endoplasmic reticulum unfolded protein response. Nat Rev Mol Cell Biol, 8(7), 519–529. https://doi.org/10.1038/nrm2199

Rudin, C. M., Durinck, S., Stawiski, E. W., Poirier, J. T., Modrusan, Z., Shames, D. S., Bergbower, E. A., Guan, Y., Shin, J., Guillory, J., Rivers, C. S., Foo, C. K., Bhatt, D., Stinson, J., Gnad, F., Haverty, P. M., Gentleman, R., Chaudhuri, S., Janakiraman, V., … Seshagiri, S. (2012). Comprehensive genomic analysis identifies SOX2 as a frequently amplified gene in small-cell lung cancer. Nat Genet, 44(10), 1111–1116. https://doi.org/10.1038/ng.2405

Rueden, C. T., Schindelin, J., Hiner, M. C., DeZonia, B. E., Walter, A. E., Arena, E. T., & Eliceiri, K. W. (2017). ImageJ2: ImageJ for the next generation of scientific image data. BMC Bioinformatics, 18(1), 529. https://doi.org/10.1186/s12859-017-1934-z

Sah, R. K., Ma, J., Bah, F. B., Xing, Z., Adlat, S., Oo, Z. M., Wang, Y., Bahadar, N., Bohio, A. A., Nagi, F. H., Feng, X., Zhang, L., & Zheng, Y. (2020). Targeted Disruption of Mouse Dip2B Leads to Abnormal Lung Development and Prenatal Lethality. Int J Mol Sci, 21(21). https://doi.org/10.3390/ijms21218223

Schrödinger, LLC. (2015). The PyMOL Molecular Graphics System, Version 1.8.

Schuhmacher, M., Grasskamp, A. T., Barahtjan, P., Wagner, N., Lombardot, B., Schuhmacher, J. S., Sala, P., Lohmann, A., Henry, I., Shevchenko, A., Coskun, U., Walter, A. M., & Nadler, A. (2020). Live-cell lipid biochemistry reveals a role of diacylglycerol side-chain composition for cellular lipid dynamics and protein affinities. Proc Natl Acad Sci U S A, 117(14), 7729–7738. https://doi.org/10.1073/pnas.1912684117

Shin, M., Ware, T. B., & Hsu, K. L. (2020). DAGL-Beta Functions as a PUFA-Specific Triacylglycerol Lipase in Macrophages. Cell Chem Biol, 27(3), 314–321 e315. https://doi.org/10.1016/j.chembiol.2020.01.005

Shubitz, L. F., Powell, D. A., Trinh, H. T., Lewis, M. L., Orbach, M. J., Frelinger, J. A., & Galgiani, J. N. (2018). Viable spores of Coccidioides posadasii Deltacps1 are required for vaccination and provide long lasting immunity. Vaccine, 36(23), 3375–3380. https://doi.org/10.1016/j.vaccine.2018.04.026

Sorger, D., & Daum, G. (2003). Triacylglycerol biosynthesis in yeast. Appl Microbiol Biotechnol, 61(4), 289–299. https://doi.org/10.1007/s00253-002-1212-4

Sorin, A., Rosas, G., & Rao, R. (1997). PMR1, a Ca2+-ATPase in yeast Golgi, has properties distinct from sarco/endoplasmic reticulum and plasma membrane calcium pumps. J Biol Chem, 272(15), 9895–9901. https://doi.org/10.1074/jbc.272.15.9895

Starr, M. L., & Fratti, R. A. (2019). The Participation of Regulatory Lipids in Vacuole Homotypic Fusion. Trends Biochem Sci, 44(6), 546–554. https://doi.org/10.1016/j.tibs.2018.12.003

Surma, M. A., Klose, C., Peng, D., Shales, M., Mrejen, C., Stefanko, A., Braberg, H., Gordon, D. E., Vorkel, D., Ejsing, C. S., Farese, R., Jr., Simons, K., Krogan, N. J., & Ernst, R. (2013). A lipid E-MAP identifies Ubx2 as a critical regulator of lipid saturation and lipid bilayer stress. Mol Cell, 51(4), 519–530. https://doi.org/10.1016/j.molcel.2013.06.014

Trivedi, O. A., Arora, P., Sridharan, V., Tickoo, R., Mohanty, D., & Gokhale, R. S. (2004). Enzymic activation and transfer of fatty acids as acyl-adenylates in mycobacteria. Nature, 428(6981), 441–445. https://doi.org/10.1038/nature02384

Tu-Sekine, B., Goldschmidt, H., & Raben, D. M. (2015). Diacylglycerol, phosphatidic acid, and their metabolic enzymes in synaptic vesicle recycling. Adv Biol Regul, 57, 147–152. https://doi.org/10.1016/j.jbior.2014.09.010

van Meer, G., Voelker, D. R., & Feigenson, G. W. (2008). Membrane lipids: where they are and how they behave. Nat Rev Mol Cell Biol, 9(2), 112–124. https://doi.org/10.1038/nrm2330

Varadi, M., Anyango, S., Deshpande, M., Nair, S., Natassia, C., Yordanova, G., Yuan, D., Stroe, O., Wood, G., Laydon, A., Zidek, A., Green, T., Tunyasuvunakool, K., Petersen, S., Jumper, J., Clancy, E., Green, R., Vora, A., Lutfi, M., … Velankar, S. (2021). AlphaFold Protein Structure Database: massively expanding the structural coverage of protein-sequence space with high-accuracy models. Nucleic Acids Res. https://doi.org/10.1093/nar/gkab1061

Veses, V., Richards, A., & Gow, N. A. (2008). Vacuoles and fungal biology. Curr Opin Microbiol, 11(6), 503–510. https://doi.org/10.1016/j.mib.2008.09.017

Vida, T. A., & Emr, S. D. (1995). A new vital stain for visualizing vacuolar membrane dynamics and endocytosis in yeast. J Cell Biol, 128(5), 779–792. https://doi.org/10.1083/jcb.128.5.779

Volmer, R., & Ron, D. (2015). Lipid-dependent regulation of the unfolded protein response. Curr Opin Cell Biol, 33, 67–73. https://doi.org/10.1016/j.ceb.2014.12.002

Volmer, R., van der Ploeg, K., & Ron, D. (2013). Membrane lipid saturation activates endoplasmic reticulum unfolded protein response transducers through their transmembrane domains. Proc Natl Acad Sci U S A, 110(12), 4628–4633. https://doi.org/10.1073/pnas.1217611110

Wang, Y., He, D., Chu, Y., Zuo, Y. S., Xu, X. W., Chen, X. L., Zhao, W. S., Zhang, Y., Yang, J., & Peng, Y. L. (2016). MoCps1 is important for conidiation, conidial morphology and virulence in Magnaporthe oryzae. Curr Genet, 62(4), 861–871. https://doi.org/10.1007/s00294-016-0593-3

Ware, T. B., Franks, C. E., Granade, M. E., Zhang, M., Kim, K. B., Park, K. S., Gahlmann, A., Harris, T. E., & Hsu, K. L. (2020). Reprogramming fatty acyl specificity of lipid kinases via C1 domain engineering. Nat Chem Biol, 16(2), 170–178. https://doi.org/10.1038/s41589-019-0445-9

Watkins, P. A. (1997). Fatty acid activation. Prog Lipid Res, 36(1), 55–83. https://doi.org/10.1016/s0163-7827(97)00004-0

Webb, B., & Sali, A. (2016). Comparative Protein Structure Modeling Using MODELLER. Curr Protoc Bioinformatics, 54, 5 6 1-5 6 37. https://doi.org/10.1002/cpbi.3

Xing, Z. K., Zhang, L. Q., Zhang, Y., Sun, X., Sun, X. L., Yu, H. L., Zheng, Y. W., He, Z. X., & Zhu, X. J. (2020). DIP2B Interacts With alpha-Tubulin to Regulate Axon Outgrowth. Front Cell Neurosci, 14, 29. https://doi.org/10.3389/fncel.2020.00029

Yang, C., & Kazanietz, M. G. (2003). Divergence and complexities in DAG signaling: looking beyond PKC. Trends Pharmacol Sci, 24(11), 602–608. https://doi.org/10.1016/j.tips.2003.09.003

Zhang, L., Mabwi, H. A., Palange, N. J., Jia, R., Ma, J., Bah, F. B., Sah, R. K., Li, D., Wang, D., Bah, F. B., Togo, J., Jin, H., Ban, L., Feng, X., & Zheng, Y. (2015). Expression Patterns and Potential Biological Roles of Dip2a. PLoS One, 10(11), e0143284. https://doi.org/10.1371/journal.pone.0143284

