## Supplementary Figures with legends for "DIP2 is a unique regulator of diacylglycerol lipid homeostasis in eukaryotes"

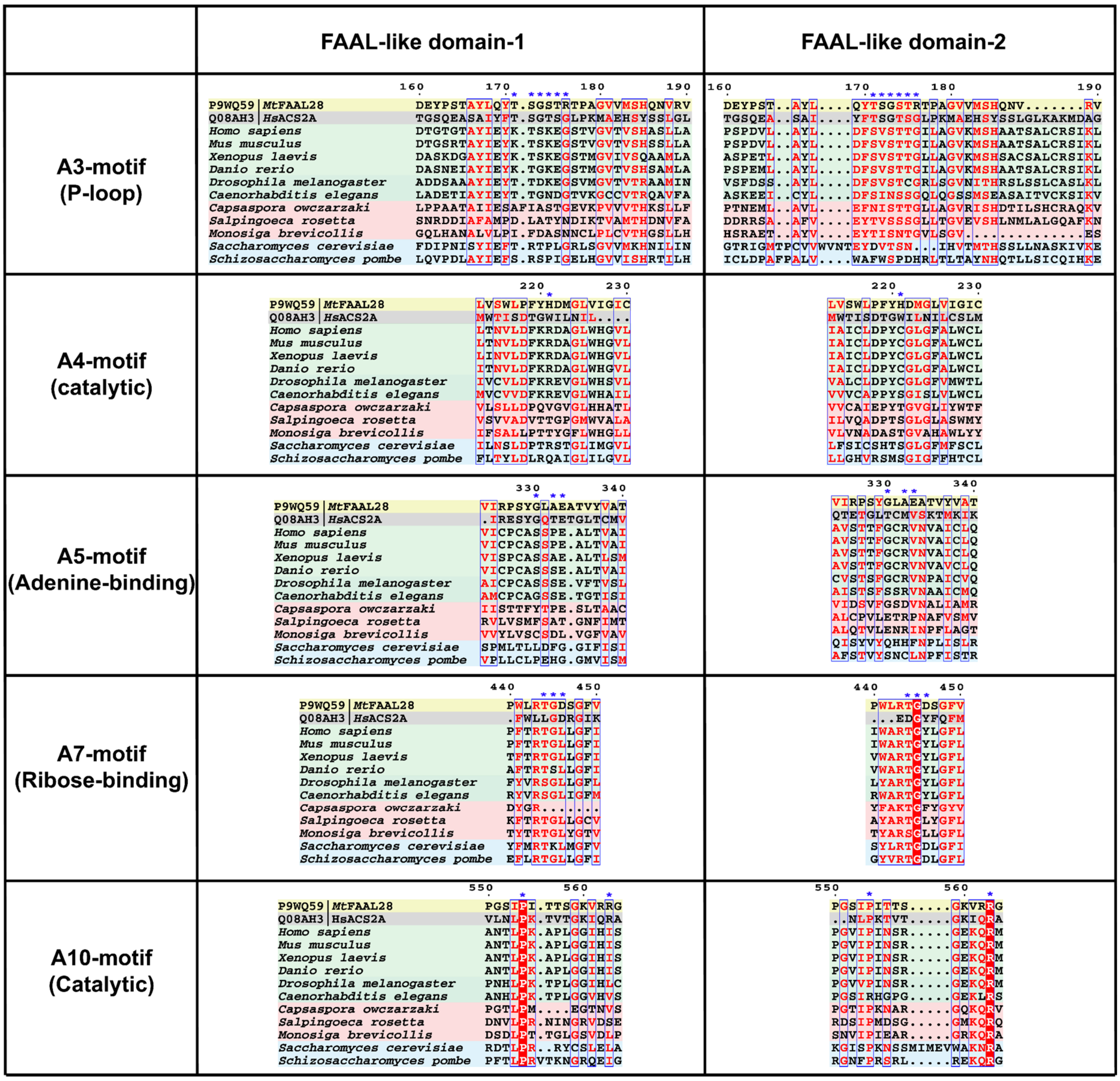
FAAL-like domain-1
FAAL-like domain-2
A3-motif (P-loop)
A4-motif (catalytic)
A5-motif (Adenine-binding)
A7-motif (Ribose-binding)
A10-motif (Catalytic)
P9WQ59 | MtFAAL28
Q08AH3 | HsACS2A
Homo sapiens
Mus musculus
Xenopus laevis
Danio rerio
Drosophila melanogaster
Caenorhabditis elegans
Capsaspora owczarzaki
Salpingoeca rosetta
Monosiga brevicollis
Saccharomyces cerevisiae
Schizosaccharomyces pombe

**Supplementary Fig. 1. Conservation and diversification of ANL superfamily motifs in FLD1 and FLD2 of DIP2.** Some of the important motifs that are conserved across the ANL superfamily such as A3 (P-loop; important for binding phosphates), A4 (Catalytic; facilitating a nucleophilic substitution on acyl-AMP), A5 (Adenine-binding; stacks against the adenine ring), A7 (Ribose-binding; tethers the hydroxyls moieties of ribose in ATP) and A10 (Catalytic; important for acyl-AMP formation) are shown. The sequence alignment shows that all these important motifs show systematic changes that are unique to each FLD and different for fungal and animal FLDs. The crucial residues of the motifs are marked with blue star sign (*). Representative FAAL, FACL, animal FLDs, choanoflagellate FLDs and fungal FLDs are highlighted with yellow, grey, green, red and blue background colors, respectively.

A

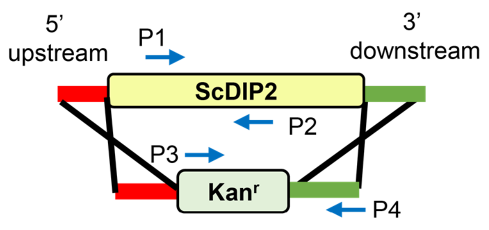


B

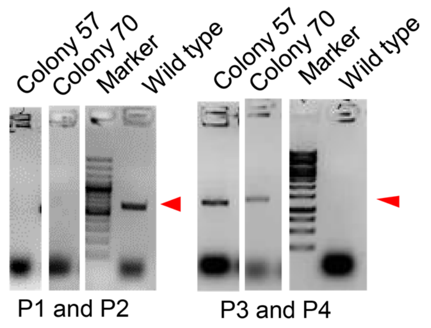


C

| GO Term | GO Term Frequency | GO Term Usage % |
| --- | --- | --- |
| Response to chemical | 24 of 183 genes | 13.11% |
| Transcription by RNA polymerase II | 23 of 183 genes | 12.57% |
| Lipid metabolic process | 19 of 183 genes | 10.38% |
| Mitotic cell cycle | 16 of 183 genes | 8.74% |
| Protein phosphorylation | 14 of 183 genes | 7.65% |
| Mitochondrion organization | 12 of 183 genes | 6.56% |
| Transmembrane transport | 12 of 183 genes | 6.56% |
| Ion transport | 10 of 183 genes | 5.46% |
| Cytoplasmic translation | 10 of 183 genes | 5.46% |

D

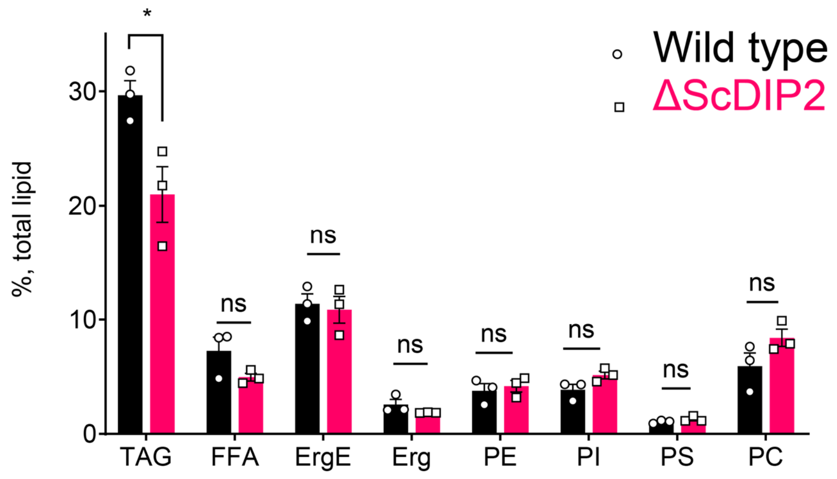

**Supplementary Fig. 2. ScDIP2 null mutant generation and metabolic radiolabeling of total lipid pool of yeast.** (**A**) Schematic for homologous recombination-based gene knock-out strategy. P1 and P2: ScDIP2 specific primers, P3 and P4: primers to confirm gene replacement. (**B**) PCR based confirmation of ScDIP2 null mutant colonies (colony 57, YSM57 and colony 70, YSM70) using above mentioned primers. PCR product bands are indicated using red arrowhead. (**C**) Genetic interactors (both positive and negative) of ScDIP2 were classified according to the Gene ontology terms. Genes involved in lipid metabolism process are among the highest interacting partners. GO terms comprise up to 5% of total interactors are shown in the table (further description in *Materials and methods* section). (**D**) Radio-TLC based quantification of major membrane lipids show no significant change between wild type and $\Delta$ScDIP2 cells. Neutral lipid, TAG shows moderate but significant depletion in $\Delta$ScDIP2 cells. All data are represented by the mean ± SEM of at least three independent experiments. *$p < 0.05$; ns= not significant. DAG, Diacylglycerol; TAG, Triacylglycerol; Erg, Ergosterol; ErgE, Ergosterol esters; FFA, Free fatty acids; PC, Phosphotidylcholine; PE, Phosphotidylethanolamine; PI, Phosphatidylinositol; PS, Phosphatidylserine.

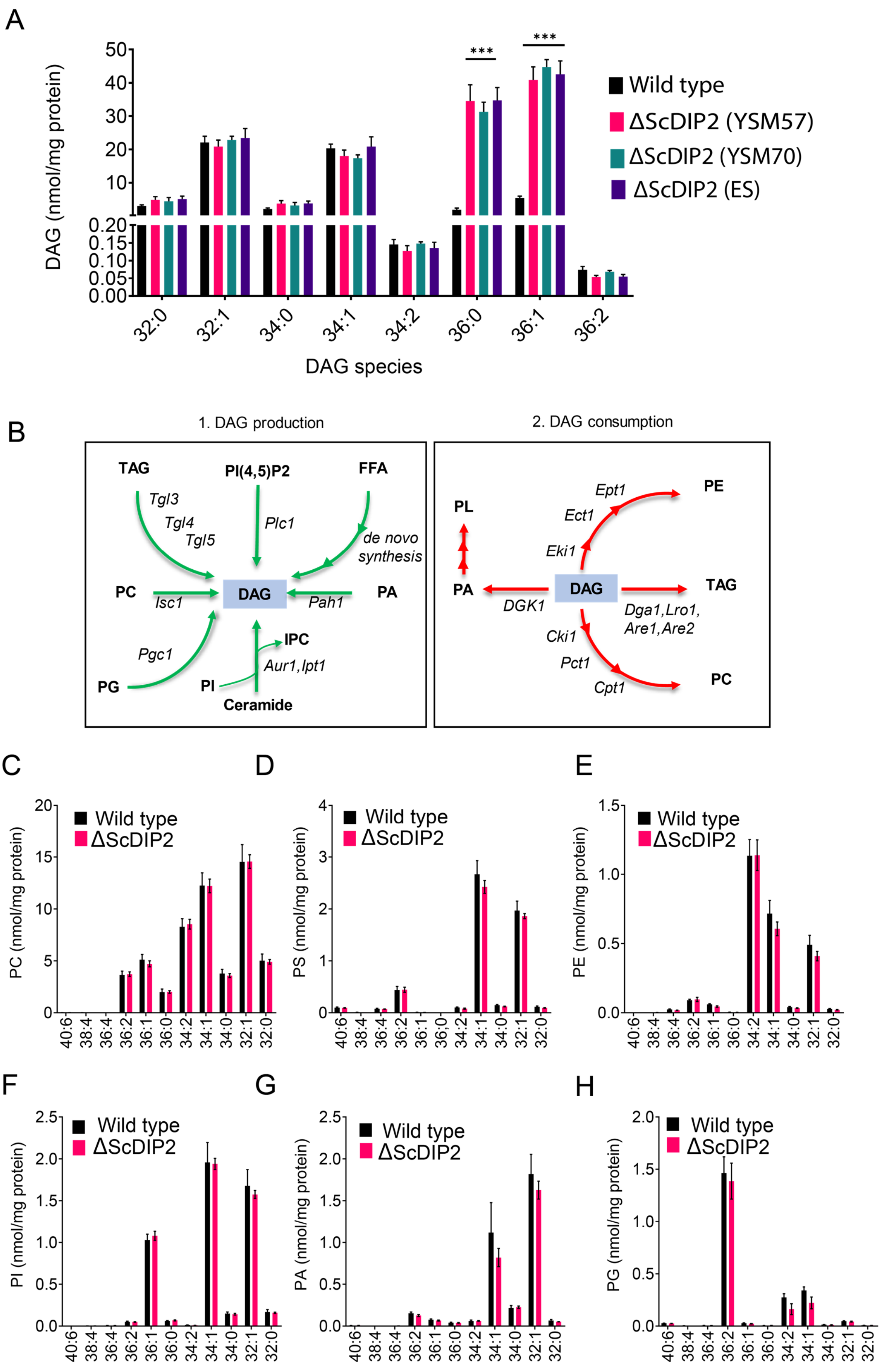
A
DAG (nmol/mg protein)
DAG species
Wild type
ΔScDIP2 (YSM57)
ΔScDIP2 (YSM70)
ΔScDIP2 (ES)
***
***
B
1. DAG production
2. DAG consumption
TAG
PI(4,5)P2
FFA
Tgl3
Tgl4
Tgl5
Plc1
de novo synthesis
PC
Isc1
DAG
Pah1
PA
Pgc1
PG
PI
IPC
Aur1,Ipt1
Ceramide
PL
PE
Ept1
Ect1
Eki1
DGK1
Dga1,Lro1, Are1,Are2
Cki1
Pct1
Cpt1
C
PC (nmol/mg protein)
D
PS (nmol/mg protein)
E
PE (nmol/mg protein)
F
PI (nmol/mg protein)
G
PA (nmol/mg protein)
H
PG (nmol/mg protein)
Wild type
ΔScDIP2

1211

**Supplementary Fig. 3. LC-MS based lipid profiling of yeast strains.** (**A**) The accumulation of DAGs is seen in multiple strains of mutant such as the mutant obtained from EUROSCARF (ES; strain Y01869) along with the two mutant colonies generated in the laboratory (ΔScDIP2 YSM57 and YSM70: in-house knock-out lines; YSM57 has been used for all other experiments in the current work) as shown in the LC-MS data. All data are represented as mean ± SEM. ***$p < 0.001$; unpaired, two-tailed student's t-test ($n = 6$). (**B**) A schematic showing multiple routes for DAG synthesis and utilization. The multiple arrowheads denote multi-step processes. The corresponding enzyme names are written on the arrows in italics while the lipid metabolites are shown in bold fonts. (**C**-**H**) Major phospholipids quantified using LC-MS showed no significant change in ΔScDIP2 cells while compared with wild type cells. Specific lipid lengths (combined number of carbons in two acyl chains) are indicated on x-axis. Data are represented as mean ± SEM ($n=4$). PA, Phosphatidic acid; PG, Phosphatidylglycerol; IPC, Inositolphosphoryl-ceramides; PI(4,5)P2, Phosphatidylinositol 4,5-bisphosphate.

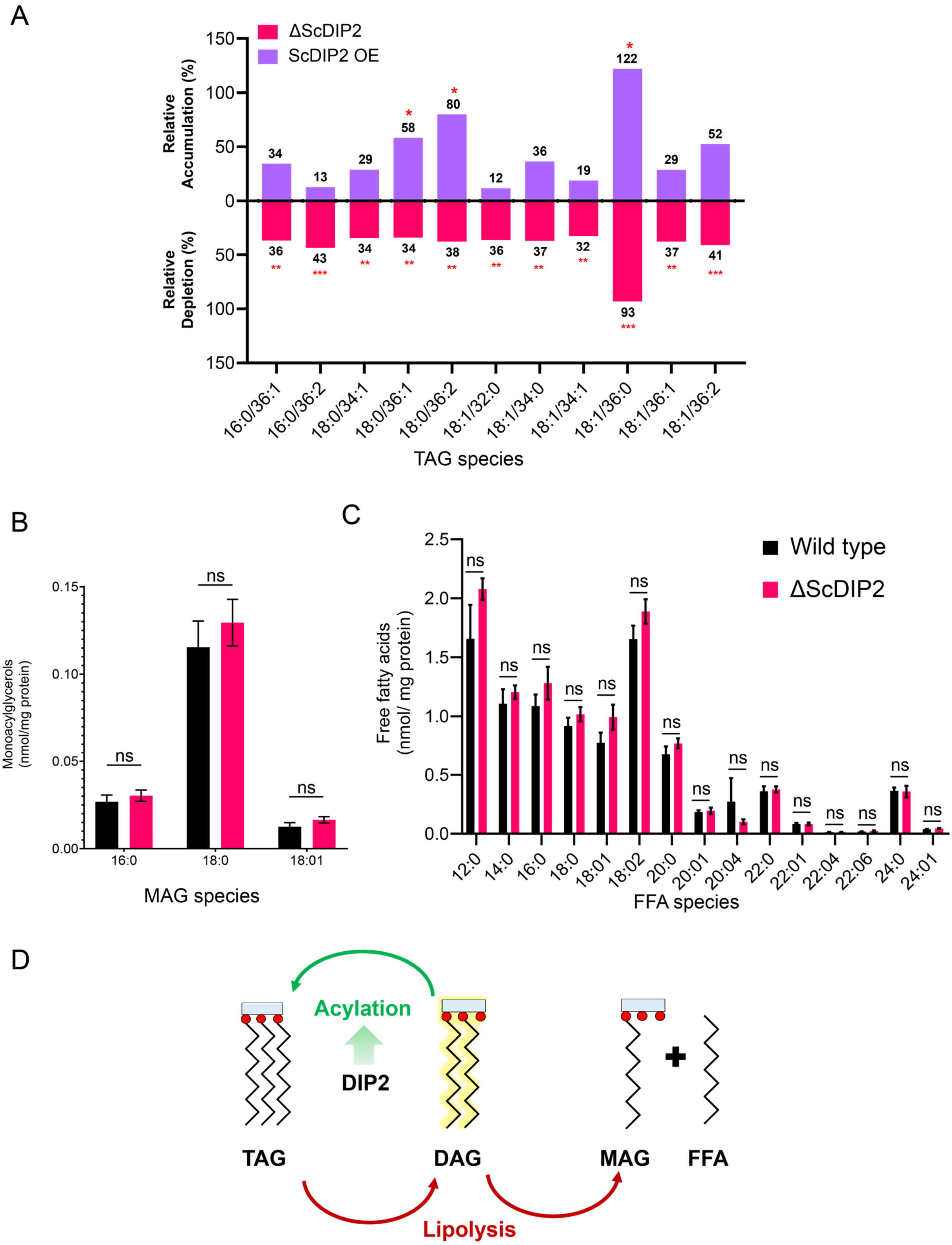
A
ΔScDIP2
ScDIP2 OE
Relative Accumulation (%)
Relative Depletion (%)
TAG species
16:0/36:1
16:0/36:2
18:0/34:1
18:0/36:1
18:0/36:2
18:1/32:0
18:1/34:0
18:1/34:1
18:1/36:0
18:1/36:1
18:1/36:2
B
Monoacylglycerols (nmol/mg protein)
MAG species
16:0
18:0
18:01
C
Free fatty acids (nmol/ mg protein)
Wild type
ΔScDIP2
FFA species
D
Acylation
DIP2
TAG
DAG
MAG
FFA
Lipolysis

**Supplementary Fig. 4. LC-MS based quantification TAG and its lipolysis products from yeast strains.** (**A**) Depletion and accumulation TAG upon deletion and overexpression of ScDIP2, respectively. Relative percent changes are shown numerically, normalized to respective wild type TAG levels (n=6; unpaired, two-tailed student's t-test; *$p < 0.05$; **$p < 0.01$; ***$p < 0.001$; ****$p < 0.0001$; ns= not significant). (**B** and **C**) Lipidomic analysis of monoacylglycerol (MAG) and free fatty acid (FFA) showed no significant changes in $\Delta$ScDIP2 cells compared with wild type (n=6; unpaired, two-tailed student's t-test; ns= not significant). (**D**) A schematic showing TAG as a metabolic source as well as the sink for the DAG pool. DAGs can be directly converted to TAGs via acylation reaction, while TAG lipolysis can also result in the formation DAG, which further breaks down into MAG and FFA.  Data suggests that the loss of DIP2 results in insufficient acylation of DAG to produce TAGs.

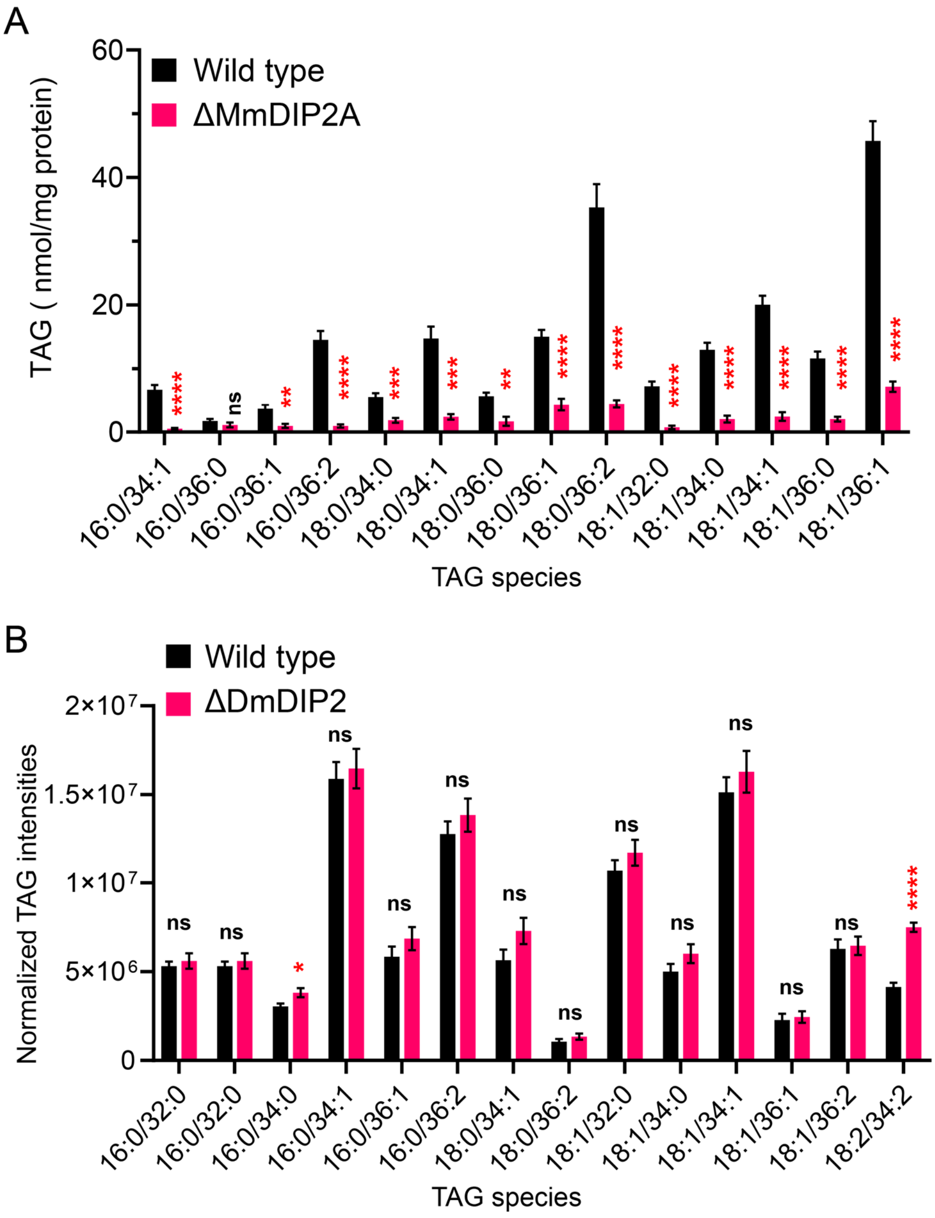
A
Wild type
ΔMmDIP2A
TAG ( nmol/mg protein)
60
40
20
0
****
ns
**
****
***
***
**
****
****
****
****
****
****
****
16:0/34:1
16:0/36:0
16:0/36:1
16:0/36:2
18:0/34:0
18:0/34:1
18:0/36:0
18:0/36:1
18:0/36:2
18:1/32:0
18:1/34:0
18:1/34:1
18:1/36:0
18:1/36:1
TAG species
B
Wild type
ΔDmDIP2
Normalized TAG intensities
2×10⁷
1.5×10⁷
1×10⁷
5×10⁶
0
ns
ns
*
ns
ns
ns
ns
ns
ns
ns
ns
ns
ns
****
16:0/32:0
16:0/32:0
16:0/34:0
16:0/34:1
16:0/36:1
16:0/36:2
18:0/34:1
18:0/36:2
18:1/32:0
18:1/34:0
18:1/34:1
18:1/36:1
18:1/36:2
18:2/34:2
TAG species

**Supplementary Fig. 5. LC-MS based TAG quantification from DIP2 knock-out mouse embryonic stem cells and *Drosophila*.** (**A**) LC-MS based quantification of TAGs from mouse embryonic stem cells (mES) show that multiple TAGs have depleted in mutant, which is consistent with the multiple DAG species that accumulated in mutant as compared to wild type. (**B**) TAGs from of 5-day old adult *Drosophila* DIP2 mutant show no significant change as compared to wild type. Unlike yeast and mES, the lipids of Drosophila were isolated from the whole body which can neutralize the chances of observing any tissue specific accumulation of TAG. Data are represented as mean ± SEM ($n > 5$; unpaired, two-tailed student's t-test, *$p < 0.05$; **$p < 0.01$; ***$p < 0.001$; ****$p < 0.0001$; ns= not significant).

A

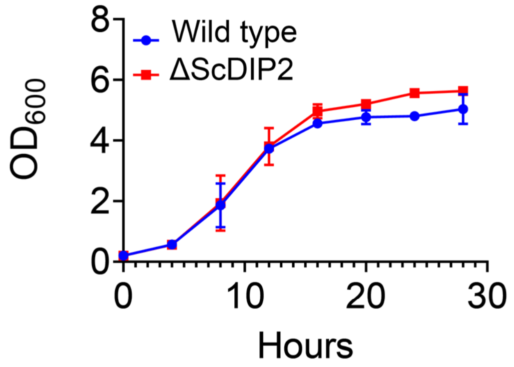


B

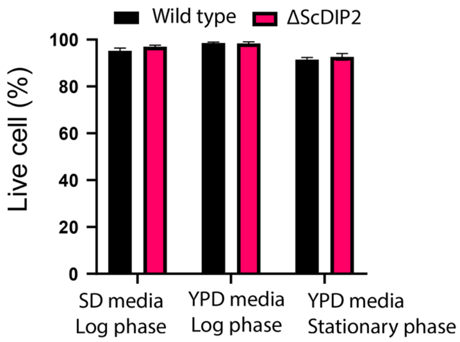


C

| Stress type | Reagent/condition | Representative data | Observation |
|---|---|---|---|
| **Plasma membrane stress** | SDS 0.005% | Wild type / ΔScDIP2 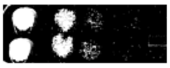 | Same as Wt |
| | Ethanol | Wild type / ΔScDIP2 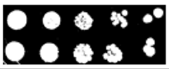 | Same as Wt |
| **Cell wall stress** | Congo red (100µg/ml) | Wild type / ΔScDIP2 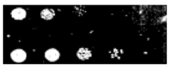 | Resistant |
| | Calcofluor white (100µg/ml) | Wild type / ΔScDIP2 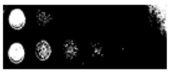 | Resistant |
| **Cold stress** | 18°C | Wild type / ΔScDIP2 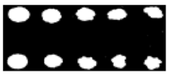 | Same as Wt |
| **Heat stress** | 37°C and 39°C | Wild type / ΔScDIP2 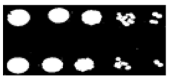 | Same as Wt |
| **Oxidative stress** | $H_2O_2$ (5mM) | Wild type / ΔScDIP2 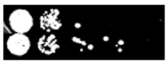 | Resistant |
| **Osmotic stress** | YPD + 1.2 M NaCl | Wild type / ΔScDIP2 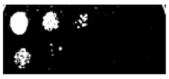 | Sensitive |
| **Starvation stress** | Nutrient depleted media | Wild type / ΔScDIP2 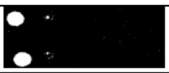 | Same as Wt |
| **Inositol auxotrophy** | SD media without inositol | Wild type / ΔScDIP2 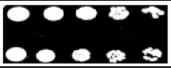 | Same as Wt |

D

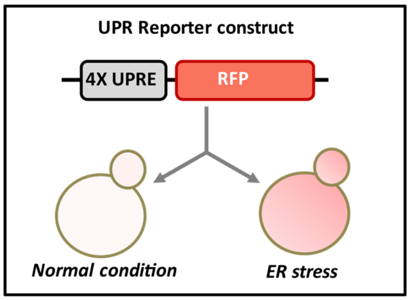


E

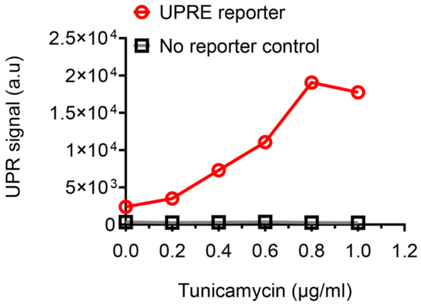


F

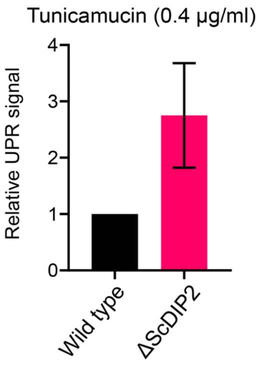

**Supplementary Fig. 6. Growth phenotype and stress responses by ΔScDIP2 cells.** (**A** and **B**) Deletion of ScDIP2 shows no growth defect as demonstrated using the growth curve and cell viability assay. Growth curve of wild type and ΔScDIP2 strain was generated at 30°C, in SD media (n=3) and cell viability assay was performed in different growth phases and growth media (n=300, triplicate). (**C**) Stress phenotype screening using spot dilution assay is shown. Growth of ΔScDIP2 cells were compared with wild type cells in indicated stress conditions. Experiments were repeated for a minimum of three times and representative data are shown. (**D**) A graphical representation of UPRE reporter assay is shown. Cells harboring 4xUPRE-RFP (mCherry) plasmid express a basal level of RFP in normal condition while, any stimuli that activates UPR pathway, will amplify RFP expression. **(E)** Fluorescence intensity of RFP was measured using FACS, which shows increase in signal intensity with increasing tunicamycin concentration in log phase yeast cells harboring the reporter plasmid. (**F**) The signal intensity from the reporter plasmid in ΔScDIP2 cells when treated with tunicamycin at concentration of 0.4 µg/ml is shown and compared to the signal output from wild type cells (mean ± SD; n=3).

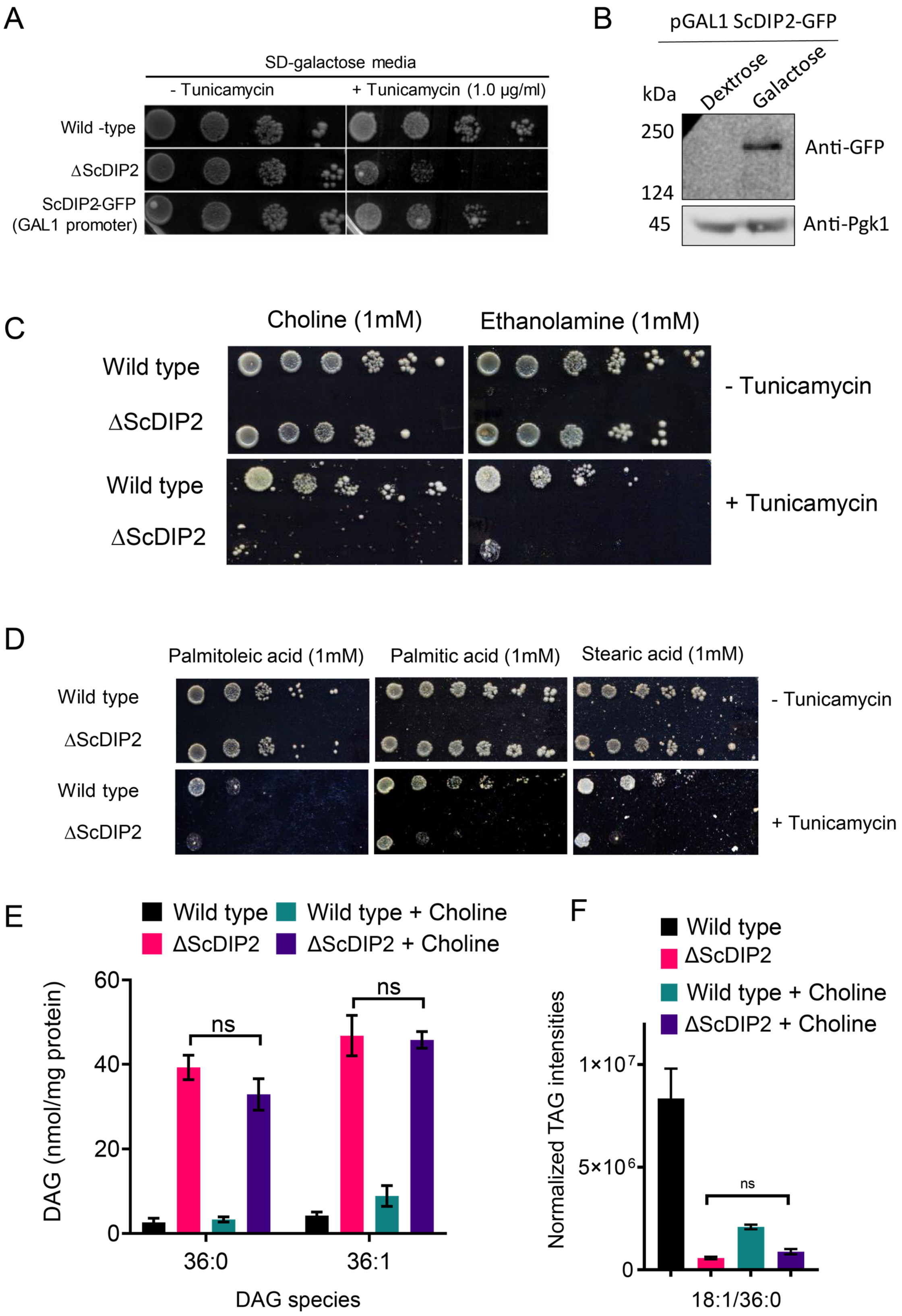
A
SD-galactose media
- Tunicamycin
+ Tunicamycin (1.0 µg/ml)
Wild -type
ΔScDIP2
ScDIP2-GFP
(GAL1 promoter)
B
pGAL1 ScDIP2-GFP
kDa
Dextrose
Galactose
250
124
45
Anti-GFP
Anti-Pgk1
C
Choline (1mM)
Ethanolamine (1mM)
Wild type
ΔScDIP2
Wild type
ΔScDIP2
- Tunicamycin
+ Tunicamycin
D
Palmitoleic acid (1mM)
Palmitic acid (1mM)
Stearic acid (1mM)
Wild type
ΔScDIP2
Wild type
ΔScDIP2
- Tunicamycin
+ Tunicamycin
E
Wild type
Wild type + Choline
ΔScDIP2
ΔScDIP2 + Choline
DAG (nmol/mg protein)
60
40
20
0
ns
ns
36:0
36:1
DAG species
F
Wild type
ΔScDIP2
Wild type + Choline
ΔScDIP2 + Choline
Normalized TAG intensities
1×10⁷
5×10⁶
0
ns
18:1/36:0

**Supplementary Fig. 7. Genetic complementation and chemical supplementation assay.** (**A**) Genetic complementation of tunicamycin-mediated ER stress sensitivity in ΔScDIP2 with galactose-inducible (GAL1 promoter) ScDIP2-GFP expression. (**B**) Western blot image is showing the full-length ScDIP2-GFP expression in galactose-supplemented growth media. Pgk1 is used as the loading control. (**C**) Screening for ER stress rescue by supplementing media with different phospholipid biosynthesis precursors like choline and ethanolamine. (**D**) Screening for ER stress rescue by supplementing with physiologically abundant fatty acids**,** indicated in the images. (**E** and **F**) No changes in selective DAG and TAG levels were observed when ΔScDIP2 cells were supplemented with choline, which is known to redirect DAG pool to phosphatidylcholine synthesis. Data are represented as mean ± SEM (n > 5; unpaired, two-tailed student's t-test; ns= not significant).

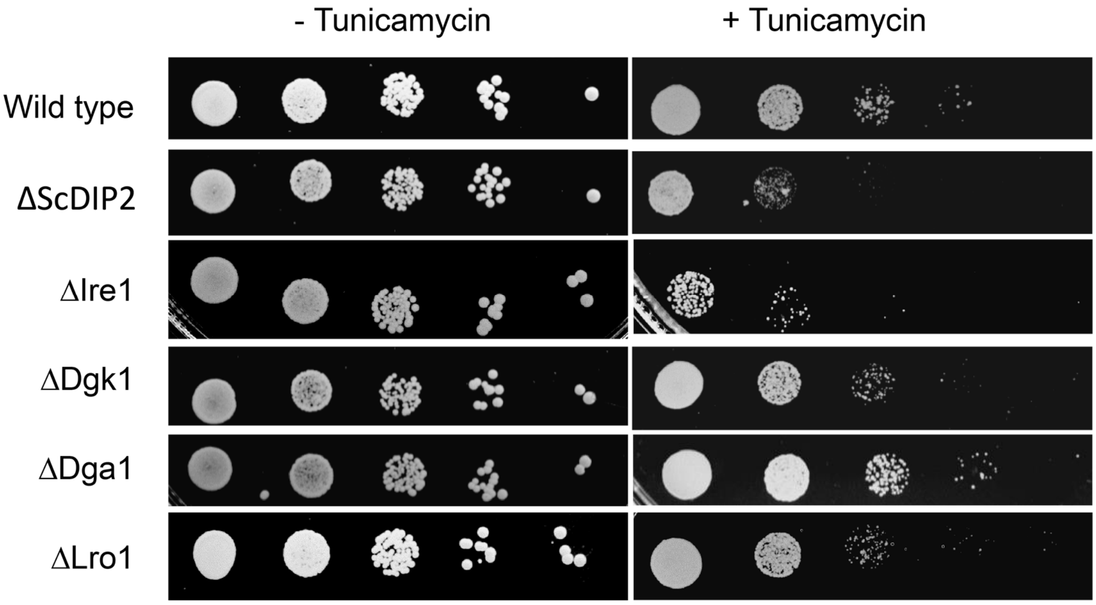

- Tunicamycin
+ Tunicamycin
Wild type
ΔScDIP2
ΔIre1
ΔDgk1
ΔDga1
ΔLro1

**Supplementary Fig. 8. ER stress sensitivity assay with null mutants of bulk DAG metabolizing enzymes.** The null mutant strains for canonical DAG metabolizing enzymes, viz. ΔDga1, Δ Lro1, ΔDgk1 (procured from EUROSCARF collection center, Germany) show less or no sensitivity to tunicamycin treatment unlike ΔScDIP2 strain (n=3, Tunicamycin concentration= 1.0 µg/ml). Null mutant of Ire1 (ΔIre1), the sensor protein of ER stress, serves as a positive control in this assay.

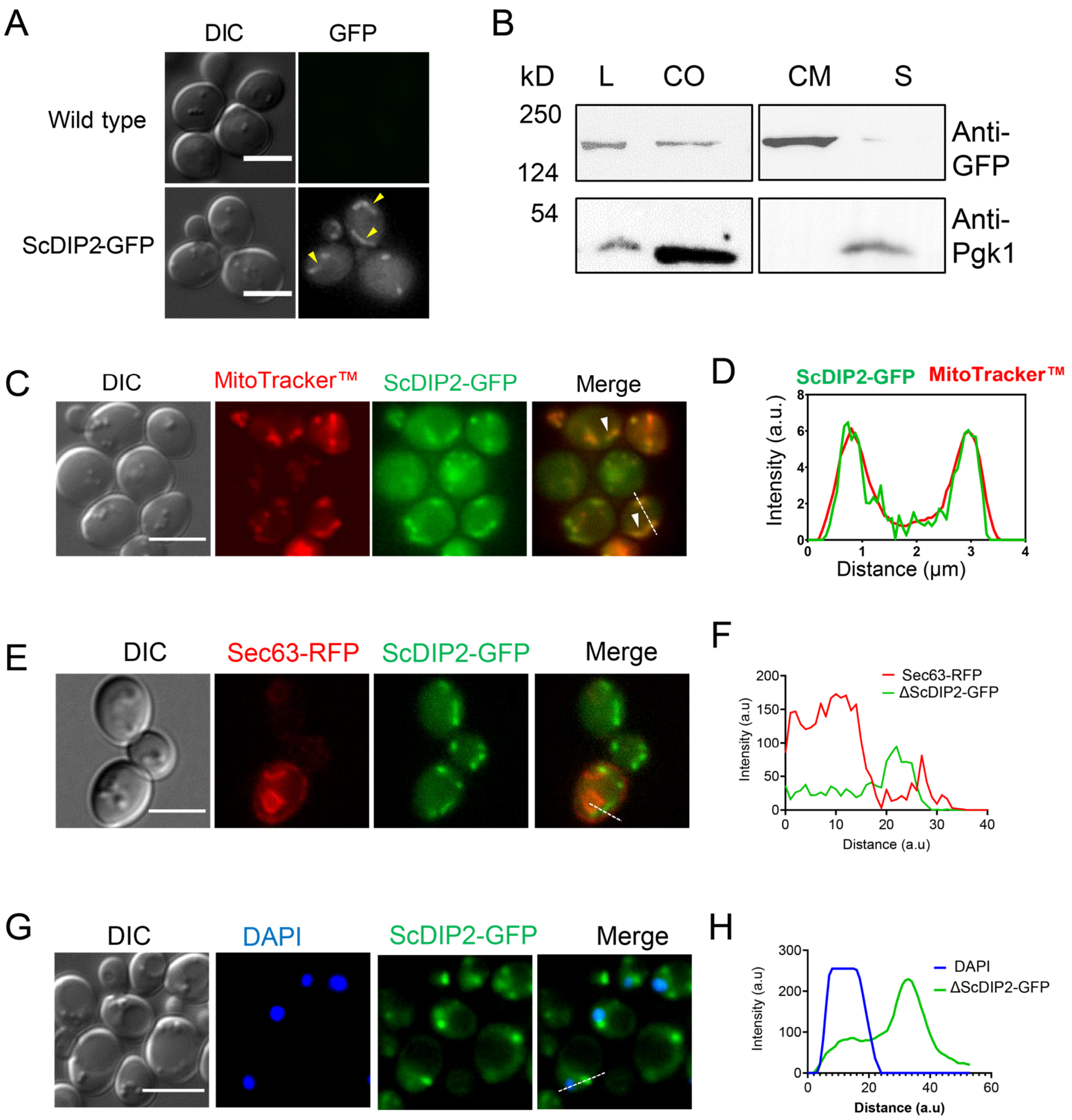
A
DIC
GFP
Wild type
ScDIP2-GFP
B
kD
L
CO
CM
S
250
124
54
Anti-GFP
Anti-Pgk1
C
DIC
MitoTracker™
ScDIP2-GFP
Merge
D
ScDIP2-GFP
MitoTracker™
Intensity (a.u.)
Distance (µm)
E
DIC
Sec63-RFP
ScDIP2-GFP
Merge
F
Sec63-RFP
ΔScDIP2-GFP
Intensity (a.u)
Distance (a.u)
G
DIC
DAPI
ScDIP2-GFP
Merge
H
DAPI
ΔScDIP2-GFP
Intensity (a.u)
Distance (a.u)

**Supplementary Fig. 9. Subcellular localization study of ScDIP2.** (**A**) ScDIP2-GFP knock-in cells shows punctate and tubular localization pattern (indicated by yellow arrow heads). Wild type cells shown under GFP channel serves as negative control. Scale bar= 5μm**.** (**B**) Western blot of the fractionated lysate from ScDIP2-GFP knock-in cells. ScDIP2-GFP was probed using anti-GFP antibody and Pgk1 is a control cytosolic protein. L= total lysate, CO= crude organelle lysate (devoid of cell wall debris and nuclei), CM= crude membrane fraction, S= membrane-less supernatant. The data shows the association of ScDIP2-GFP with crude membrane fraction. (**C**) ScDIP2-GFP is associated with mitochondria as seen from the colocalization of GFP and MitoTracker$^{TM}$ intensities. Scale bar= 5μm**.** (**D**) The line scan of signal intensity along the white line shown in (C). (**E** and **G**) ScDIP2-GFP shows no significant subcellular association with ER marker, Sec63-RFP and nuclear stain, DAPI. Scale bar= 5μm. (**F** and **H**) Line scan analysis along the dotted line to show the no significant colocation of fluorescence signal from indicated channels shown in (E) and (G).

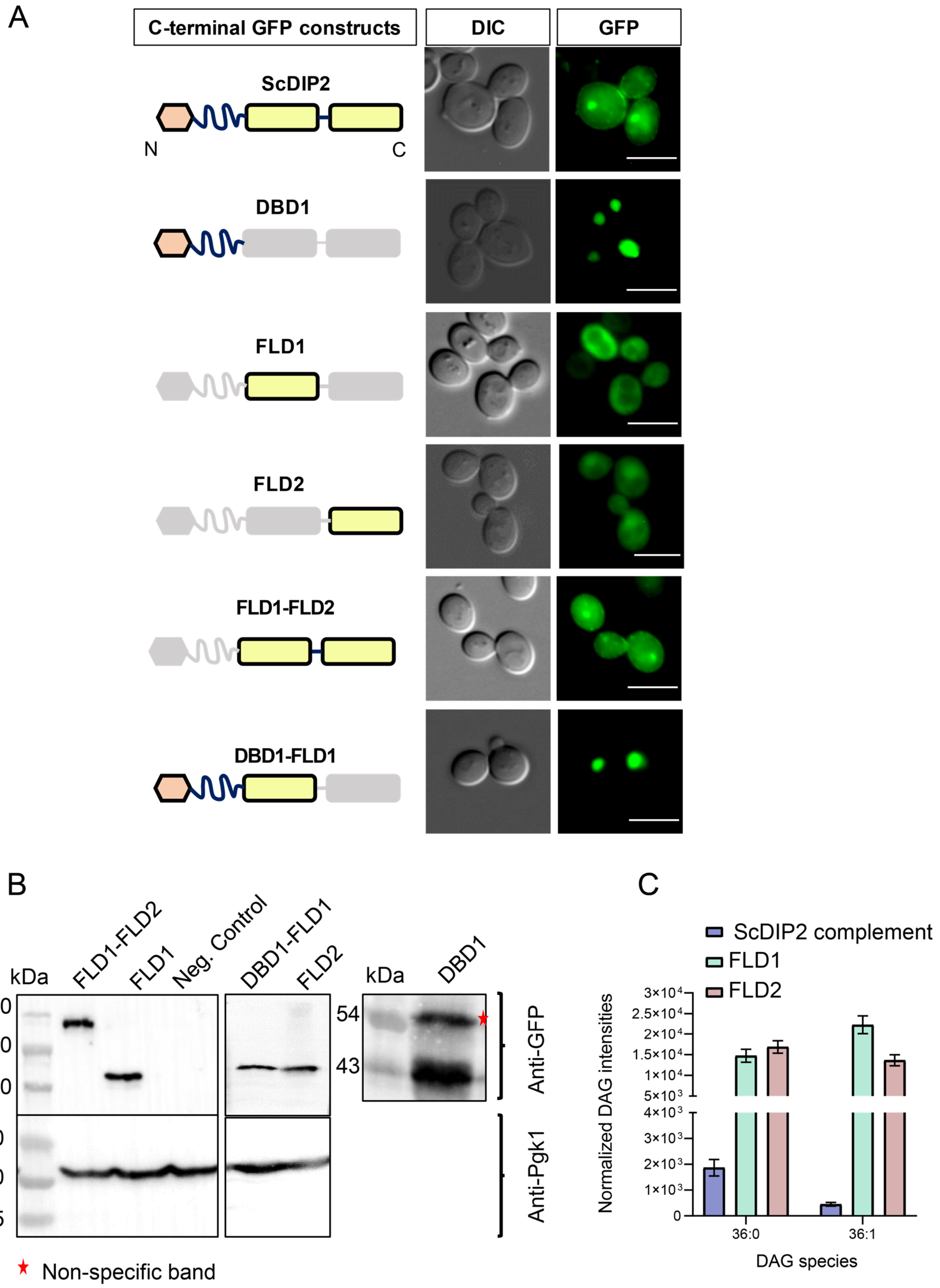
A
C-terminal GFP constructs
DIC
GFP
ScDIP2
N
C
DBD1
FLD1
FLD2
FLD1-FLD2
DBD1-FLD1
B
kDa
FLD1-FLD2
FLD1
Neg. Control
DBD1-FLD1
FLD2
250
150
100
50
40
35
kDa
DBD1
54
43
Anti-GFP
Anti-Pgk1
Non-specific band
C
ScDIP2 complement
FLD1
FLD2
Normalized DAG intensities
DAG species
36:0
36:1

**Supplementary Fig. 10. Expression validation and lipidomics of ScDIP2 domain-truncated constructs.** (**A** and **B**) Representative fluorescence microscopy images and western blots of various C-terminal GFP-tagged domain-truncated constructs of ScDIP2 (expressed under GAL1 promoter). Deleted part of domains are shown in gray color. The constructs were used for the ER stress rescue assay by complementing $\Delta$ScDIP2 strains and subsequent lipidomic analysis. Scale bar= 5µm. (**C**) LC-MS quantification of C36:0 and C36:1 DAGs from $\Delta$ScDIP2 cells complemented with individual FLD domains (FLD1 and FLD2) and full length ScDIP2 expressed under GAL1 promoter. Unlike the full-length ScDIP2, individual FLD expression is not able to restore selective DAG level. Data are represented as mean $\pm$ SEM ($n > 5$; unpaired, two-tailed student’s t-test).

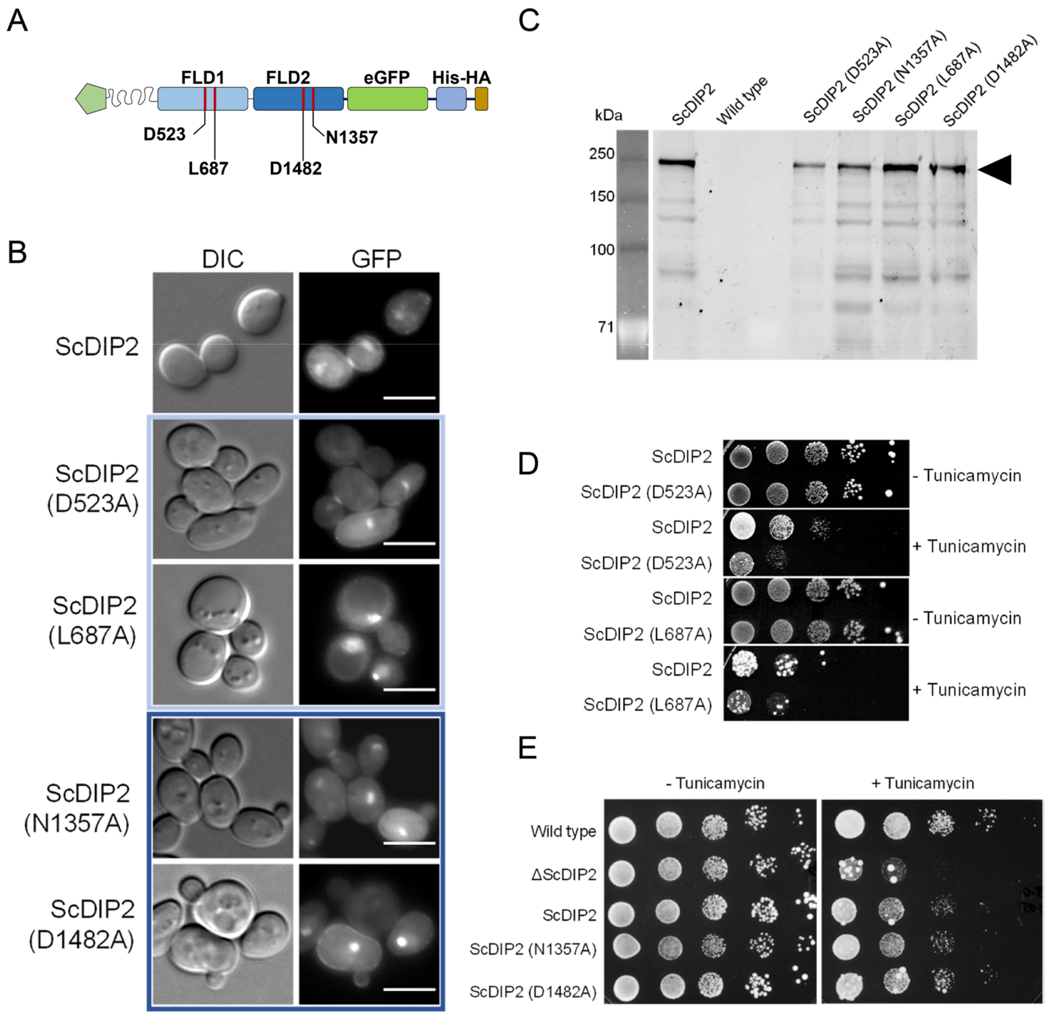
A
FLD1
FLD2
eGFP
His-HA
D523
L687
D1482
N1357
B
DIC
GFP
ScDIP2
ScDIP2 (D523A)
ScDIP2 (L687A)
ScDIP2 (N1357A)
ScDIP2 (D1482A)
C
kDa
250
150
100
71
ScDIP2
Wild type
ScDIP2 (D523A)
ScDIP2 (N1357A)
ScDIP2 (L687A)
ScDIP2 (D1482A)
D
ScDIP2
ScDIP2 (D523A)
- Tunicamycin
+ Tunicamycin
ScDIP2 (L687A)
E
- Tunicamycin
+ Tunicamycin
Wild type
ΔScDIP2
ScDIP2
ScDIP2 (N1357A)
ScDIP2 (D1482A)

**Supplementary Fig. 11. Probing the catalytic role of ScDIP2 via mutational analysis of FLDs.** (**A**) A schematic to depict the residue names and positions that were used for mutational analysis. (**B**) Fluorescence microscopy images of wild type and mutant ScDIP2 variants. Scale bar = 5 µm. (**C**) Expression of the full-length mutant ScDIP2 variants checked in SDS-PAGE through GFP fluorescence channel (at 488 nm). (**D** and **E**) Representative images of tunicamycin-induced ER stress assay to screen loss of function in mutant ScDIP2 variants (n=3, Tunicamycin concentration= 1.0 µg/ml).
