## Supplementary Tables 1-4 for "DIP2 is a unique regulator of diacylglycerol lipid homeostasis in eukaryotes"

**Supplementary Table 1:**

| | Organism | Mutation | Phenotype | Disease/Pathogenesis | Reference |
|---|---|---|---|---|---|
| *Fungi* | *Cochliobolus heterostrophus* | ΔDIP2 | Reduced or loss of virulence | Southern leaf blight | (Lu et al., 2003) |
| | *Cochliobolus victoriae* | | Reduced or loss of virulence | Victoria blight | |
| | *Gibberella zeae* | | Reduced or loss of virulence | Fusarium head blight, | |
| | *Magnaporthe oryzae* | ΔDIP2 | Loss of Virulence | Rice blast/blight | (Wang et al., 2016) |
| | *Coccidioides posadasii* | ΔDIP2 | Loss of Virulence | Valley fever | (Narra et al., 2016; Shubitz et al., 2018) |
| *Animals* | *Caenorhabditis elegans* | ΔDIP2 | Increased ectopic neurite sprouting, branching and axon regeneration | - | (Noblett et al., 2019) |
| | *Drosophila melanogaster* | ΔDIP2 | Axon branching and guidance defect in mushroom-body neurons | - | (Nitta et al., 2017) |
| | *Mus musculus* | ΔDIP2a | Defective dendritic spine morphogenesis and reduced synaptic transmission | Autism-like behaviour | (Ma et al., 2019) |
| | | | Diet dependent growth defects | - | (Kinatukara et al., 2020) |
| | | ΔDIP2b | Excessive axonal outgrowth, Reduced synaptic transmission | - | (Xing et al., 2020) |
| | | | Defective lung formation, Peri-natal lethality | - | (Sah et al., 2020) |
| | *Homo sapiens (RKO cell line)* | ΔDIP2c | Induction of epithelial-mesenchymal transition and enhanced cell motility | | (Larsson et al., 2017) |
| | *Homo sapiens* | DIP2a [#] | - | Developmental dyslexia | (Kong et al., 2016; Poelmans et al., 2009) |
| | | | - | Autism | (Egger et al., 2014; Iossifov et al., 2012) |
| | | DIP2b [#] | - | Coronary artery disease | (Gong et al., 2018) |
| | | DIP2c [#] | - | Cancers | (Jiao et al., 2012; Rudin et al., 2012) |

#= SNP (single-nucleotide polymorphism), de novo frameshift/ nonsense variant, deletions

**Supplementary Table 2:** List of yeast strains.

| **Strain ID** | **Genotype** | **Source** |
|---|---|---|
| BY4741 | MATa his3Δ1 leu2Δ0 met15Δ0 ura3Δ0 | (Brachmann et al., 1998) |
| YSM57, YSM70 (ΔScDip2) | MATa; ura3Δ0; leu2Δ0; his3Δ1; met15Δ0; YOR093c::kanMX4 | This study |
| Y01869 (ΔScDip2) | MATa; ura3Δ0; leu2Δ0; his3Δ1; met15Δ0; YOR093c::kanMX4 | EUROSCARF |
| Y01907 (ΔIre1) | BY4741; MATa; ura3Δ0; leu2Δ0; his3Δ1; met15Δ0; YHR079c::kanMX4 | EUROSCARF |
| Y02501 (ΔDga1) | BY4741; MATa; ura3Δ0; leu2Δ0; his3Δ1; met15Δ0; YOR245c::kanMX4 | EUROSCARF |
| Y05383 (ΔLro1) | BY4741; MATa; ura3Δ0; leu2Δ0; his3Δ1; met15Δ0; YNR008w::kanMX4 | EUROSCARF |
| Y01608 (ΔDgk1) | BY4741; MATa; ura3Δ0; leu2Δ0; his3Δ1; met15Δ0; YOR311c::kanMX4 | EUROSCARF |
| YSM101 | MATa; ura3Δ0; leu2Δ0; his3Δ1; met15Δ0; YOR093c:: GFP(S65T)-KanMX6 | This study |

**Supplementary Table 3:** List of Primers.

| Name | Primer type | Primer sequences |
|---|---|---|
| CMR2-P1-FP | Forward | CAGCGTACGTTGCGTCTTA |
| CMR2-P2-RP | Reverse | CAATGACGATTGCAAAGAAGC |
| Kan-P3-FP | Forward | GTGACGACTGAATCCGGTG |
| CMR2-P4-RP | Reverse | CGCAGCTTCTCAGTGTTAC |
| CMR2-Del-5'-FP | Forward | GTGTTAGGGTGATTCAGTTCTGTGTAAAAGCGTGTGGCATTGAGTTACTCCAATGCGTACGCTGCAGGTCGAC |
| CMR2-Del-5'-RP | Reverse | CACAACTCAGTAGCATCCAATAGTCATGACAAATTTACTGTACTTGGATGTGTTAATCGATGAATTCGAGCTCG |
| CMR2-pGAL1- FP | Forward | GTCAAGGAGAAAAAACCCCATTAATTAAATGGATTTTTCTATTCCTCCTACC |
| CMR2-pGAL1-RP | Reverse | AGTACAGGTTTTCTCCGGACTCGAGAATATTGTCCTTTTCATAATCTGATAATAAATAAATGGAAATATTTTCAC |
| ScDMAP1-PGAL1-Gib-RP | Reverse | GAAGTACAGGTTTTCTCCGGACTCGAGTGCTGAGTCTCCACTGCTAGTATTTTCG |
| ScF2-pGAL1-Gib-FP | Forward | GTCAAGGAGAAAAAACCCCATTAATTAAATGGTTAAACCAAAACTTGCCCTACAATGC |
| ScF1F2-pGAL1-Gib-FP | Forward | GTCAAGGAGAAAAAACCCCATTAATTAAATGACGGATTCTTTACCGCTAATTTTACG |
| ScF1-pGAL1-Gib-RP | Reverse | GAAGTACAGGTTTTCTCCGGACTCGAGGAGATCATTGTTTAAAAACTTCTTCTCTACCGTG |
| PCMR2T-pPM90-FP | Forward | ACCTTGCATGCTCTCTACAATTAGCTTGTCTTTTC |
| PCMR2T-pPM90-RP | Reverse | AGCTCGGTACCAAGTTCGTTAAGGAAGAAACAAGAT |

| | | |
|---|---|---|
| ScDip2-GFP-KanMX6-KI-FP | Forward | GGCATAAACTATGGTGAAAATATTTCCATTTATTTATTATCAGATTATGAAAAGGACAATATTGGTGGCAGTAAAGGAGAAGAACTTTTCACTGG |
| ScDip2-GFP-KanMX6-KI-RP | Reverse | CCACAACTCAGTAGCATCCAATAGTCATGACAAATTTACTGTACTTGGATGTGATCGATGAATTCGAGCTCGTTTAAACTGGATGG |
| ScDIP2-D523A-FP | Forward | CCCATGTTAACGTTATTGGCTTTTGGTGGTATCTTTATATCTATAAGAGATCA |
| ScDIP2-D523A-RP | Reverse | CTTATAGATATAAAGATACCACCAAAAGCCAATAACGTTAACATGGGAGAATA |
| ScDIP2-L687A-FP | Forward | CAAATACCTACTTTATGAGAACCAAGGCTATGGGGTTTGTTCATAACGGAAAGAT |
| ScDIP2-L687A-RP | Reverse | CCGTTATGAACAAACCCCATAGCCTTGGTTCTCATAAAGTAGGTATTTGCAGGAC |
| ScDIP2-N1357A-FP | Forward | GTTATGTCTATCAACATCACTTCGCTCCGCTTATATCATTAAGGTCGTATCTG |
| ScDIP2-N1357A-RP | Reverse | CGACCTTAATGATATAAGCGGAGCGAAGTGATGTTGATAGACATAACTTATTT |
| ScDIP2-D1482A-FP | Forward | TTAAGCTATTTGAGAACTGGTGCTCTGGGCTTTATCAAAAACGTAAGTTG |
| ScDIP2-D1482A-RP | Reverse | CGTTTTTGATAAAGCCCAGAGCACCAGTTCTCAAATAGCTTAAAGTGTTAT |

**Supplementary Table 4:** List of plasmids.

| Plasmid ID | Description | Reference |
|---|---|---|
| pYSM8 | pPPM90-CEN-URA3-YcpLac33 | Dr. Palani Murugan (CSIR-CCMB) |
| pYSM1 | pLE124 | Dr. Palani Murugan (CSIR-CCMB) |
| pYSM6 | pFA6a-GFP(S65T)-KanMX6 | Dr. Venkat Chalamcharla (CSIR-CCMB) |
| pYSM5 | pGAL1-MCS-TEV-GFP-8XHis-2XHA | This study |

| | | |
|---|---|---|
| pYSM7 | pPPM90-PROMOTER-ScDIP2-TERMINATOR | This study |
| pYSM10 | pGAL1-ScDIP2-TEV-GFP-8XHis-2XHA | This study |
| ScDIp2 Domain truncated constructs | pGAL1-ScDBD1-TEV-GFP-8XHis-2XHA | This study |
| | pGAL1-ScDBD1-TEV-GFP-8XHis-2XHA | This study |
| | pGAL1-ScFLD1-TEV-GFP-8XHis-2XHA | This study |
| | pGAL1-ScFLD2-TEV-GFP-8XHis-2XHA | This study |
| | pGAL1-ScFLD1FLD2-TEV-GFP-8XHis-2XHA | This study |
| | pGAL1-ScDBD1FLD1-TEV-GFP-8XHis-2XHA | This study |
| pPM47 (4xUPRE-RFP) | pPM47 (UPR-RFP CEN/ARS URA3) | (Merksamer et al., 2008) |
| pSM1960 | pRS426-SEC63-mRFP | (Metzger et al., 2008) |
